# Genetic differentiation of water penny beetles may be associated with the formation process of the ancient Lake Biwa

**DOI:** 10.1101/2022.11.15.516684

**Authors:** Natsuko Ito Kondo, Machiko Nishino, Makoto Uenishi, Kanako Ishikawa, Nobuyoshi Nakajima, Noriko Takamura

## Abstract

**Background:** Water flow is one of several factors that determine the habitats of aquatic insects. The conversions of stagnant (lentic) and running (lotic) water are physiologically and ecologically constrained. In general, aquatic insect species only occur in one of these environments. Though *Eubrianax ramicornis* (Coleoptera: Psephenidae) usually resides in rivers, however, it is also known to occur near the shores of Lake Biwa in Japan. Here, we investigated the genetic differences between the two Eubrianax species of Lake Biwa and its inflowing rivers, and other regions of Japan for the first time. Lake Biwa is an ancient lake and about four million years old. Though over 60 endemic species were described from Lake Biwa, they are mostly fish and molluscs, with only two species of aquatic insects. There are few reports of endemic aquatic insects in other ancient lakes, too. Discussion of the endemism of Eubrianax and its promoting factors is expected to provide insights into the evolution of aquatic insects in ancient lakes.

**Methods:** We surveyed and morphologically identified psephenid larvae in Lake Biwa and its inflowing rivers during the summers of 2017 and 2018 (Hayashi, 2009: Hayashi & Sota, 2009). We conducted phylogenetic analyses of two regions of COI and estimated the ages of population divergence based on fossil records of *Eubrianax* beetles in Lake Biwa as well as those of previously reported populations in other regions. We also analyzed two COI regions of the *E. ramicornis* haplotypes in Lake Biwa.

**Results:** Four psephenid species were identified among the samples collected in Lake Biwa and its inflowing rivers. Of these, *E. ramicornis* was collected only at the Lake Biwa shore whereas its congeneric *E. granicollis* was collected only from the rivers inflowing the lake. The phylogenetic analyses showed that *E. ramicornis* of Lake Biwa were clearly differentiated from those in previously reported regions and this divergence was estimated to have occurred 1.40 MYA. At that time, there was no lake at the location of present-day Lake Biwa. It is thought that rivers and their dammed flood plains to the east, with gravel and sand supplied by the rising Suzuka Mountains further to the east. Such an environments is preferable for *E. ramicornis* and they were thought to have become isolated from the surrounding rivers during the formation of Lake Biwa. On the contrary, there was no genetic differentiation between the *E. granicollis* population at the Lake Biwa inflow rivers and those elsewhere. Hence, *E. granicollis* might have migrated to the Lake Biwa watershed later than *E. ramicornis* and maintained gene flow over a wide area. The genetic differentiation of *E. ramicornis* reported here for the first time suggests that a species with a wide distribution may rather be evolving endemic to Lake Biwa. Reports on the endemism of aquatic insects in ancient lakes are scarce. This study is expected to bring new perspectives to the study of insect evolution in ancient lakes.

## Introduction

Aquatic insects are highly diverse and > 100,000 species have been described worldwide. They may account for 60% of all known freshwater organisms (Dijkstra, Monaghan & Pauls, 2014). Geological history involves connections between continents and islands by glacial periods, volcanic and orogenic activity, and lake formation and is considered an important contributing factor in insect evolution. Aquatic insects have adapted to widely differing environments through ecological adaptation and allopatric speciation (Dijkstra, Monaghan & Pauls, 2014). Terrestrial waters account for only 1% of the earth’s total surface area. Nevertheless, they are highly diverse in terms of their salinity, temperature, dissolved oxygen (DO) content, bottom sediment texture and composition, and geographical features (Koroiva & Pepinelli, 2019). Fluidity also greatly influences aquatic insect adaptation. In lotic environments, water flows like a river whereas in lentic environments such as puddles, ponds, and lakes, it stands or only slightly moves. In general, aquatic insects are restricted to one habitat or the other at species or certain taxonomic groups (Ribera & Vogler, 2000; Dijkstra, Monaghan & Pauls, 2014). Lotic environments are usually oxygen-rich because of constant agitation of the water. However, the organisms residing there have had to evolve morphological and behavioral traits such as flattening or clinging to avoid being swept away by currents. In lentic environments, the water is calm but its DO concentration is low even during normal conditions. Hence, organisms dwelling in lentic areas have had to evolve traits enabling them to absorb oxygen efficiently (Gullan & Cranston, 1994). Organisms that have physiologically and morphologically adapted to the water flow in their physical environments cannot readily switch between lotic and lentic habitats.

Differences in the stability and connectivity of lentic and lotic water habitats may affect population genetic structure (Ribera & Vogler, 2000; Ribera, Bilton & Vogler, 2003; Abellan, Millan & Ribera, 2009). In the previous studies, it was assumed that lotic environments tend to be more stable and continuous than lentic environments. Therefore, lotic species need not migrate over long distances and are only slightly characterized by the gene flow associated with populations migration and dispersal. On the other hand, lentic habitats are discontinuous and often unstable because their waters occasionally dry up. Therefore, lentic species have high mobility and dispersal ability, maintain gene flow over a broad area, and are comparatively less genetically structured. The foregoing statements comprise the “habitat-stability-hypothesis” (Ribera & Vogler, 2000; Ribera 2008). Results supporting this theory have been reported for aquatic beetles, crustaceans, and mollusks (Marten, Brandle & Brandle, 2006; Abellan, Millan & Ribera, 2009; Arribas et al., 2012). However, other studies have also reported discordant results. It was proposed that physical water flow alone does not influence population genetic structure or differentiation. The latter might also be affected by migration barriers and habitat characteristics such as stability and persistence in long-term, historical, and geological backgrounds (Letsch, Gottsberger & Ware, 2016; Saito & Tojo, 2016; Tojo et al., 2017; Takenaka et al., 2021).

Over 200 species of water penny beetle (Psephenidae) have been reported and are globally distributed. Most of them produce larvae that live in freshwater (Hayashi, 2009) and are considered to indicate water quality. Nevertheless, it is difficult to identify them based on their larval morphology alone. For this reason, their utility as indicator species is limited in practical field work (Hayashi, 2009). Psephenid larvae generally occur in lotic habitats such as rivers but only rarely in lentic habitats, especially large-scale lakes (Hayashi, 2009; Lee, Yang & Sato, 2001; Jung, Jach & Bae, 2020). However, a few of *Eubrianax* species inhabit in large lakes (Murvosh, 1992; Nishino, 1992; Hayashi, 2009; Tsutsumi, 2017). Twenty-three *Eubrianax* species and subspecies are listed worldwide. Of these, six occur on the main island of Japan while another six live on the Ryukyu Islands (Hayashi, 2009; Hayashi, Song & Sota, 2012). *E. ramicornis* is widely distributed and lives in the lotic environments and may be found in its reservoirs with small inflowing rivers of the Korean Peninsula, the Tsushima, Goto, and Oki Islands, and all main islands of Japan except Hokkaido (Hayashi, 2009). *E. ramicornis* also inhabits in Lake Biwa, the largest lake in Japan. Similarly, the North American *Eubrianax edwardsii* usually inhabits in rivers but also occurs in the few large lakes of California (Murvosh, 1992). On the other hand, the genetic differentiation between riverine and large lake populations of either species is unknown.

Lake Biwa is located in the center of Honshu Island and is the ancient lake. This lake originated from Paleo-Lake Biwa ~4 MYA and the present-day lake basin began to form about 0.43 MYA (Satoguchi 2012; Satoguchi 2020). Ancient lakes have long-term stability and might have served as refugia during the past glacial periods. Their substantial area, volume, and depth provide diverse horizontal and vertical habitats and promote the evolution of species assemblages with rapid adaptive dispersal (Cristescu et al., 2010). The faunal taxa that have diversified in these environments include cichlids in the African great lakes (Seehausen, 2002; Verheyen et al. 2003; Seehausen, 2006), cottoids and amphipods in Lake Baikal (Kontula, Kirilchik & Väinölä, 2003; Gurkov et al., 2019), and amphipods and gastropods in Lake Ohrid (Budzakoska-Gjoreska et al., 2014; Stelbrink et al., 2016; Grabowski, Wysocka & Mamos, 2017). Mechanisms that promote rapid habitat differentiation and adaptation within ancient lakes include geographic isolation and hybridization caused by the emergence and disappearance of satellite lakes and small rivers in response to water level changes and lake drying (Cristescu et al., 2010).

The flora and fauna of Lake Biwa and its adjacent areas are as diverse as those in other ancient lakes. Over 3,100 different taxa have been reported for Lake Biwa (Nishino, 2020). About 150 macrofaunal taxa were reported exclusively for Lake Biwa, and 64 of these are endemic (Nishino, 2020; Ichise et al. 2021; Sawada & Nakano 2021). For fish, forty-six indigenous taxa were present and of these, 17 were reported as endemic. Nevertheless, the proportions of endemic species are lower in Lake Biwa than other ancient lakes. By contrast, > 80% of all species are endemic to Lakes Tanganyika and Baikal (Martens, 1997; Tabata et al., 2016; Maehata, 2020). Lake Biwa may have migrated from the southeast to the north through repeated appearances and disappearances of paleo-lakes and finally emerged at its present location. The newly formed lake basin then expanded and deepened (Yokoyama, 1995; Satoguchi, 2020). These successive basin migrations created various habitats at different times and led to the evolution of diverse organisms in response to global climate change. Tabata et al. (2016) reported that the multiple occurrences of fish gene flow between Lake Biwa and Paleo-Lake Biwa and the rivers and lakes in other regions differed from the events that occurred in typical ancient lakes with numerous endemic species. The authors speculated that Lake Biwa may have been a reservoir or source for the surrounding rivers that dried up because of past climate change.

Several studies of genetic diversity and phylogenetic analyses of Lake Biwa endemics of fish and shellfish have ensured their endemism (Tabata et al., 2016; Hirano et al., 2019; Miura et al., 2019; Kaneko et al., 2021). However, only two aquatic insect species have been reported as endemic for Lake Biwa. One possible reason for only a few reports of endemic aquatic insect species in Lake Biwa is that most adult insects have a far greater dispersal ability than fish or mollusks. Hence, it may be difficult for aquatic insects to be genetically differentiate. Moreover, it may also simply be due to the fact that there are fewer studies of genetic differentiation in aquatic insects than in fish and mollusks. The endemic caddisfly *Apatania biwaensis* (Nishimoto, 1994) is closely related to *Apatania aberrans* (Martynov) and the latter is the most common *Apatania* species on Honshu Island. *A. biwaensis* exhibits variable genitalia morphology even among the populations on the northern and eastern shores of North Lake (Nishimoto, 1994). The mayfly *Ephoron limnobium* (Ishiwata, 1996) is another endemic species of Lake Biwa. However, there is also an argument that *E. limnobium* might, in fact, be a junior synonym of *Ephoron shigae* (Sekiné et al., 2013; Tojo et al., 2017; Ishiwata & Uenishi, 2020). Nevertheless, these species have distinctly different egg morphology and it has been suggested that they may live separately in rivers (*E. shigae*) and within the lakes (*E. limnobium*) (Ishiwata, 1996; Ishiwata & Uenishi, 2020). Further research is needed to draw conclusions. Though it has not yet been described as an endemic species, a lineage of the widely distributed mayfly *Ecdyonurus yoshidae* was recently reported to be endemic to Lake Biwa (Kaneko et al., 2021). There are certainly few insect species whose morphology is clearly endemic to Lake Biwa. As in the case of *E. yoshidae*, however, further researches of species distributed over a wide area, including Lake Biwa, would be expected to reveal the possibility of the existence of genetically differentiated groups.

Few endemic species of aquatic insects have been reported in other ancient lakes as well as in Lake Biwa. Therefore, studies on the endemism of *E. ramicornis* in Lake Biwa and the factors driving it are expected to provide useful insights into the evolution of aquatic insects in ancient lakes. Genetic analyses on *E. ramicornis* have been conducted on individuals from Japanese and Korean rivers (Hayashi & Sota, 2012; Jung, Jach & Bae, 202). However, there have been no studies to verify whether there is genetic differentiation between those lotic populations and those found in Lake Biwa. In this study, we surveyed larvae of Psephenidae near the shore of Lake Biwa and its inflowing rivers. We analyzed the genetic distances and molecular phylogenetic relationships of *E. ramicornis* and its close relative *Eubrianax granicollis* collected in the survey with previously reported individuals from other regions. Based on the estimated divergence time and the distribution of haplotypes within Lake Biwa, the genetic differentiation of both species and its relation to the geological history of the Lake Biwa formation were discussed.

## Materials & Methods

### Field survey and identification

The coastal area of Lake Biwa and its major inflowing rivers were surveyed at 17 sites in 2017 and 2018 (Table 1). Surveys were conducted from August 1–4, 2017 at nine sites on the North basin lakeshore, at five sites on the inflowing river to the North basin, and at one site on the South basin lakeshore. In 2018, surveys were conducted from August 22–24 and on August 30 at ten sites and one site on the of the North and South basin lakeshores, respectively.

**Table 1.**
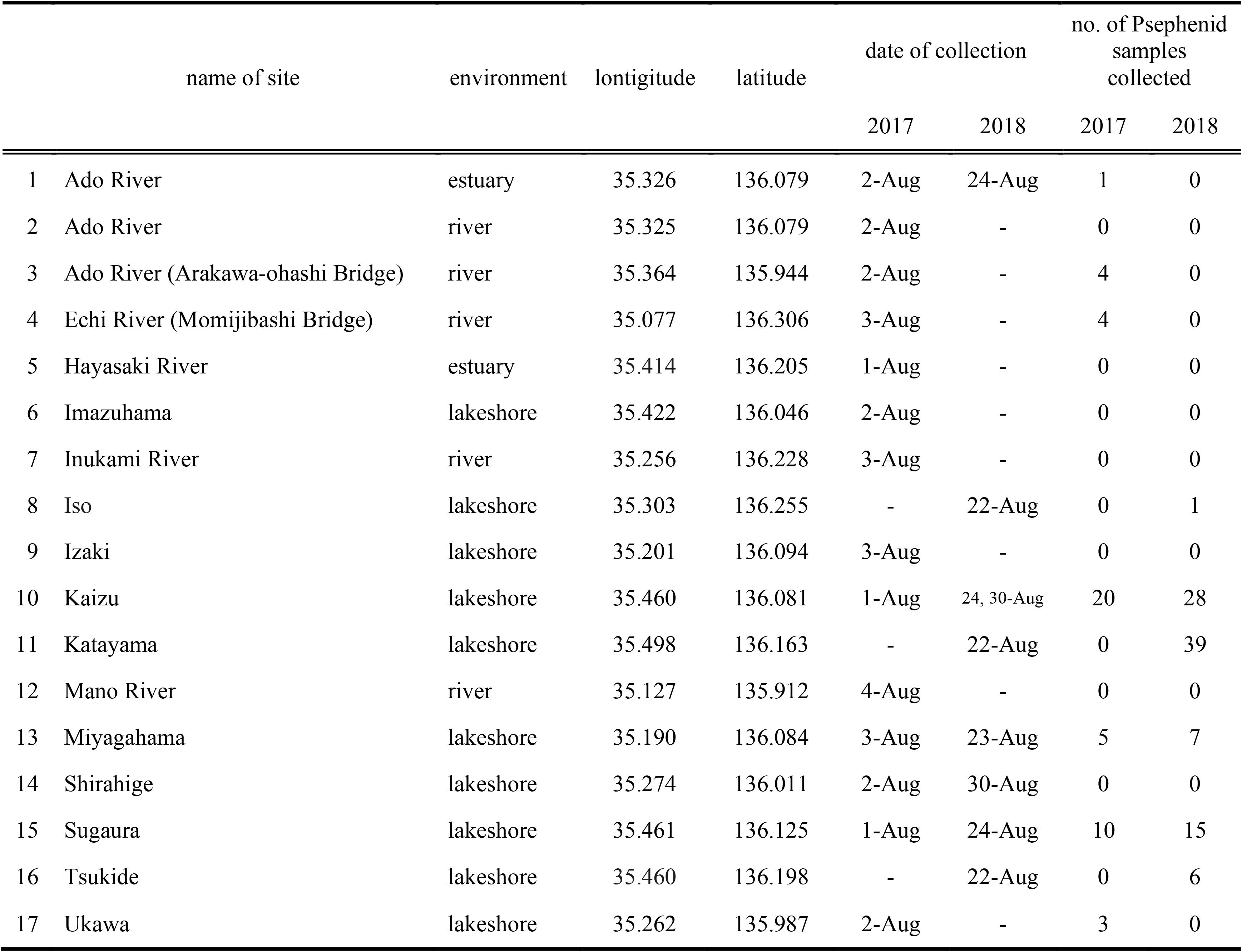
Lakeshore and inflowing river study sites of Lake Biwa.

Larval Psephenidae live on underwater rocks. Therefore, we collected rocks and gravels on the bottom of the lake and rivers at < 1 m depth. Tweezers were used to place the larvae into plastic trays and the specimens were then fixed in 99.5% (v/v) ethanol. In the laboratory, specimens were identified under a binocular and photographed according to the methodology of Hayashi (2009) and Hayashi & Sota (2008). The specimens were then returned to the ethanol for further preservation.

### DNA extraction and sequencing

DNA from the morphologically identified specimens was extracted with a DNeasy Blood and Tissue Kit (QIAGEN, Hilden, Germany). The specimens collected in 2017 were ground in 180 μL of ATL buffer with 20 μL proteinase K and incubated at 56 °C for 3 h. The specimens collected in 2018 were placed intact in ATL buffer with proteinase K, incubated overnight, and returned to the ethanol. After incubation, 200 μL ATL buffer and 200 μL of 99.5% (v/v) ethanol were added and the mixture was vortexed and purified with elution buffer according to the kit manufacturer’s directions. The final elution volume was 100 μL.

Three primer sets used to sequence the two regions of cytochrome *c* oxidase subunit I (COI) gene. LCO1490 and HCO2198 (Former et al., 1994) amplified a fragment of the region at the 5’ end of the COI gene. This region is standard for animal DNA barcoding (Herbert et al., 2003a) and is hereafter designated “LH”. The other two primer sets used to sequence were COS-Psephenid and COA-Psephenid (Hayashi & Sota, 2008) and C1-J-2195 and TL2-N-3014, respectively (Simon et al., 1994). Both sets amplified nearly the same 3’ region of COI which is hereafter designated “CT”. Most sequences already published for the *Eubrianax* beetles of Japan were in the CT region (Hayashi & Sota, 2008; Hayashi, Song & Sota, 2012). PCR was performed using GoTaq Hot Start Green Master Mix (Promega, Madison, WI, USA) under the following conditions: initial denaturation at 95 °C for 2 min, denaturation cycles at 95 °C for 45 s each, annealing at each temperature for 45 s, extension at 72 °C for 60 s, and a final extension at 72 °C for 5 min. The annealing temperature and number of PCR cycles for each primer set are shown in Table S1. The PCR products were purified with ExoSAP-IT (Applied Biosystems Inc. (ABI), Foster City, CA, USA) or the fragments of relevant size were extracted with E-gel Size Select II (Thermo Fisher Scientific, Waltham, MA, USA), sequenced with a BigDye Terminator v. 3.1 Cycle Sequencing Kit (ABI), and analyzed with ABI3730 (ABI). The sequences were then aligned with MEGA X (https://www.megasoftware.net/dload_win_gui) (Kumar et al. 2018). The accession numbers of the nucleotide sequences obtained in this study were LC572173–LC572257 and LC669409–LC669412 registered to DNA Data Bank of Japan (DDBJ; Tables S2–S4).

### Haplotypes and network analyses

There were intraspecific variations in the COI sequences of *E. ramicornis* which was the most abundant species collected in the present study. To clarify the geographical patterns of these variations in Lake Biwa, the LH and CT sequences of 30 individuals from six sites were classified into a single haplotype if they differed by a single base. Phylogenetic networks were reconstructed from these LH sequences with Network v. 10.1.0.0 (Fluxus Engineering. Ltd.; https://www.fluxus-engineering.com/sharenet.htm). The initial network was calculated by using the Median Joining (MJ) algorithm (Bandelt et al., 1995) followed by the Maximum Parsimony (MP) option (Polzin & Daneschmand, 2003) at all its default settings.

### Phylogenetic analyses

The larvae were phylogenetically analyzed using the sequences obtained in this study as well as those for *Eubrianax* spp. reported in Hayashi & Sota (2008), Hayashi, Song & Sota (2012), and Jung, Jach & Bae (2020) (Table S2). The CT regions of the COI sequences were aligned with CLUSTAL W using the default settings in MEGA X (Kumar et al., 2018) and trimmed into 720-bp fragments. A phylogenetic tree with maximum likelihood (ML) analysis was estimated using MEGA X. The divergence time was analyzed with BEAST2 v. 2.6.2 (Bouckaert et al., 2019). The inferred phylogenetic trees were graphically described with FigTree (https://github.com/rambaut/figtree/releases) (Rambaut, 2012) and organized with Adobe Illustrator 2020 in such a manner that they did not affect the results of the analysis.

To select the optimal substitution pattern model for the ML analysis, evolutionary analyses were conducted in MEGA X using 131 sequences and those of *Mataeopsephus japonicus* and *Malacopsephenoides japonicus* as outgroups. TN93 (Tamura-Nei) + G + I was selected as the model with the lowest BIC (Bayesian information criterion) value. ML analyses were performed using the selected model by bootstrapping for 1,000 iterations.

To estimate divergence time in *Eubrianax*, 133 sequences were subjected to Bayesian model analysis in BEAST2 v. 2.6.2 (Drummond & Bouckaert, 2015; Bouckaert et al., 2019). In total, 128 sequences were used in the ML tree except the haplotype CT1a sequence. Five sequences served as outgroups according to the methodology of Hayashi, Song & Sota (2012). The gamma + HKY model was used as the substitution model and the strict clock model as the clock model. All other model parameters were the default values. For calibration, divergence age priors from two fossil records were used according to the methodology of Hayashi, Song & Sota (2012). The first node was the most recent common ancestor (MRCA) of the *Eubrianax pellucidus* species group comprising *E. pellucidus, Eubrianax manakikikuse, Eubrianax amamiensis*, and *Eubrianax insularis* (Lee, Yang & Sato, 2001). The underlying fossil discovered in Tottori on Honshu Island, Japan was estimated to be an Upper Miocene Tatsumitoge member (ca. 6.5–5.5 MYA) resembling *E. pellucidus* (Hayashi & Kawakami, 2009). Hence, this fossil may be an ancestor of the *E. pellucidus* species group, and the MRCA node of the latter could be > 5.5 MYA (Hayashi, Song & Sota, 2012). The second calibration node was based on the date of divergence of the COI gene in *Plateumaris* leaf beetles estimated using the fossil record (Sota & Hayashi, 2007). Node height priors of the MRCA for *Eubrianax* spp. were acquired by using the COI sequence divergence calculated by Hayashi, Song & Sota (2012). The two calibration nodes were designated “Ep” and “E”, respectively. The node age priors for both calibration nodes were calculated by log distribution with means 6.0 MYA for node Ep and 7.11 MYA for node E and standard deviation (SD). A calibrated Yule model was used for the tree prior, and the default settings were used for all other parameters. The MCMC chain length was 10,000,000, burn-in was 100,000, and sampling was conducted every 1,000 iterations. The log file was opened with Tracer v. 1.7.1 (http://beast.community/tracer) (Rambaut et al., 2018) and it confirmed the convergence of all parameters with effective sample size (ESS) > 200. The trees file was summarized into a single tree with TreeAnnotator v. 2.6.2 (https://beast.community/treeannotator) in BEAST2 (Drummond & Bouckaert, 2015; Bouckaert et al., 2019).

### Genetic distance

The genetic distances between and within *E. ramicornis, E. brunneicornis*, and *E. granicollis* were calculated by using MEGA X (Kumar et al., 2018) with the Kimura 2-parameter (K2P) model. The analysis involved ninety-eight 720-bp nucleotide sequences of the CT region (Table S2, S3).

## Results

### Species identification

The larva of *E. granicollis, E. ramicornis, Mataeopsephus japonicus* (*Mat. japonicus*), and *Malacopsephenoides japonicus* (*Mal. japonicus*) (Table 2) were collected on the lakeshore and inflowing rivers of Lake Biwa and identified based on their morphology (Hayashi, 2009; Hayashi & Sota, 2008) and the LH and CT regions of their COI sequences. The sequence of the LH region of *Mat. japonicus* was indeterminate as its sequencing quality was substandard. A BLAST search confirmed that the sequences of the CT regions of *E. granicollis* and *E. ramicornis* were consistent with those in GenBank. The sequence of the CT region of *Mal. japonicus* had 96% identity with that of the same species in Korea. As of September 27, 2022, however, no sequences registered for *E. granicollis*, *E. ramicornis*, or *Mal. japonicus* had > 90% match in the LH region.

**Table 2.** Psephenid larvae collected in the present study. Study sites wherein no psephenids were collected are not shown.

| Site | Type | total no. | *E. ramicornis* | *E. granicollis* | *Mal. japonicus* | *Mat. japonicus* |
| --- | --- | --- | --- | --- | --- | --- |
| Momijibashi Bridge, Echi River | river | 4 | 0 | 4 | 0 | 0 |
| Arakawa-ohashi Bridge, Ado River | river | 4 | 0 | 4 | 0 | 0 |
| Ado River | estuary | 1 | 0 | 0 | 1 | 0 |
| Sugaura | lakeshore | 25 | 25 | 0 | 0 | 0 |
| Kaizu | lakeshore | 31 | 31 | 0 | 0 | 0 |
| Ukawa | lakeshore | 3 | 3 | 0 | 0 | 0 |
| Miyagahama | lakeshore | 12 | 12 | 0 | 0 | 0 |
| Iso | lakeshore | 1 | 0 | 0 | 0 | 1 |
| Tsukide | lakeshore | 6 | 6 | 0 | 0 | 0 |
| Katayama | lakeshore | 15 | 14 | 0 | 1 | 0 |

### Geographical distribution

We collected psephenid larvae at 10 of the 17 lakeshore and inflowing river sites surveyed at Lake Biwa (Table 2). The most abundant species was *E. ramicornis* and all of them were collected from six lakeshore sites on Lake Biwa. No larvae of *E. ramicornis* was collected from the inflowing rivers. Conversely, *E. granicollis* was collected from two sites in the middle reaches of the Ado and Echi Rivers flowing into Lake Biwa. However, no individual was collected from the shore of Lake Biwa. *Mat. japonicus* was collected only at the Iso on the Lake Biwa shore while *Mal. japonicus* was collected at one site on the lakeshore and one site at the estuary of the Ado River. The various species were collected sympatrically at Katayama where both *E. ramicornis* and *Mal. japonicus* were collected in 2018. In all other cases, only single species were collected at each field site.

### Phylogenetic analysis

The COI phylogenetic tree estimated with the ML model showed that the *E. ramicornis* from the shores of Lake Biwa were monophyletic with those from the other regions of Japan and in Korea (Fig. 1). Within the Japanese *E. ramicornis* clade, however, the Lake Biwa specimens diverged from those of other regions. The *E. ramicornis* from Japanese sites other than Lake Biwa were divided into two geographically unexplained groups. The clade consisting of *E. brunneicornis*, that had formerly described as a subspecies of *E. ramicornis* (Hayashi, Song & Sota, 2012), showed clear divergence from *E. ramicornis. E. granicollis* consisted of a monophyletic clade with a mixture of individuals from inflow river of Lake Biwa and those from other regions. There was no geographical trend in divergence within the clade of *E. granicollis*.

**Figure 1.**
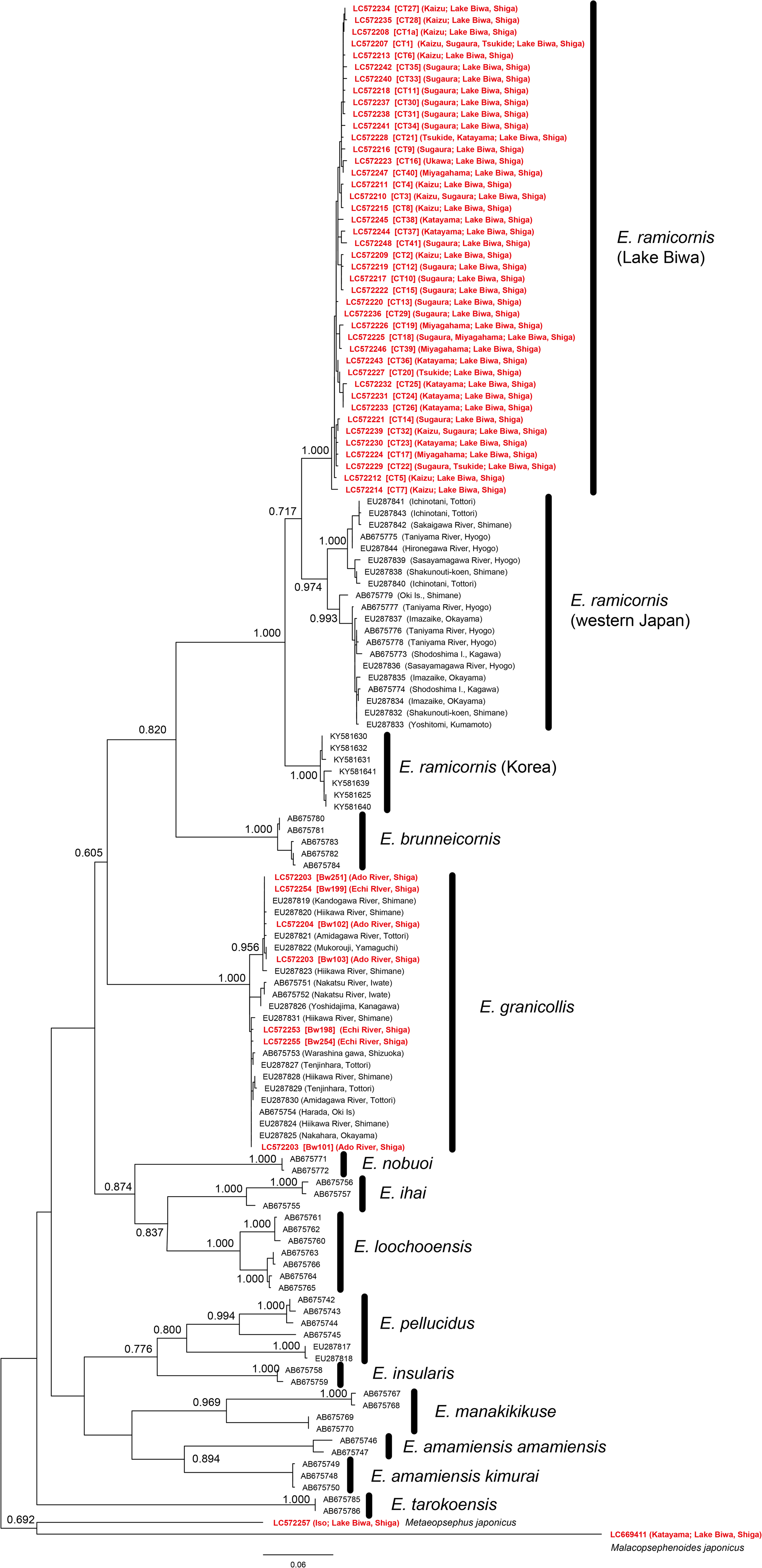
Phylogenetic tree estimated from COI sequences of *Eubrianax* by maximum likelihood model. 720-bp sequences were used. Operational taxonomic unit (OTU) names are shown in accession numbers. Specimens collected are indicated in red. Square brackets and parentheses show haplotype names/samples and collection sites, respectively. Numbers at nodes are bootstrap values > 60% after 1,000 iterations.

### Time tree

The divergence time of *Eubrianax* was estimated using two nodes as fossil-based branching ages for calibration (Fig. 2). Nodes E and Ep were the divergence times of the genus *Eubrianax* and the *E. pellucidus* species group, respectively. The mean divergence times estimated for nodes E and Ep were 7.11 MYA (median, 7.08 MYA; SD, 0.47 MYA; 95% HPD (highest posterior density); 8.02–6.22 MYA) and 5.95 MYA (median, 5.94 MYA; SD, 0.25 MYA; 95% HPD; 6.47–5.48 MYA), respectively. These estimates indicated that the median divergence time of the Lake Biwa *E. ramicornis* population from those of other regions was 1.40 MYA (95% HPD; 1.78–1.07 MYA).

**Figure 2.**
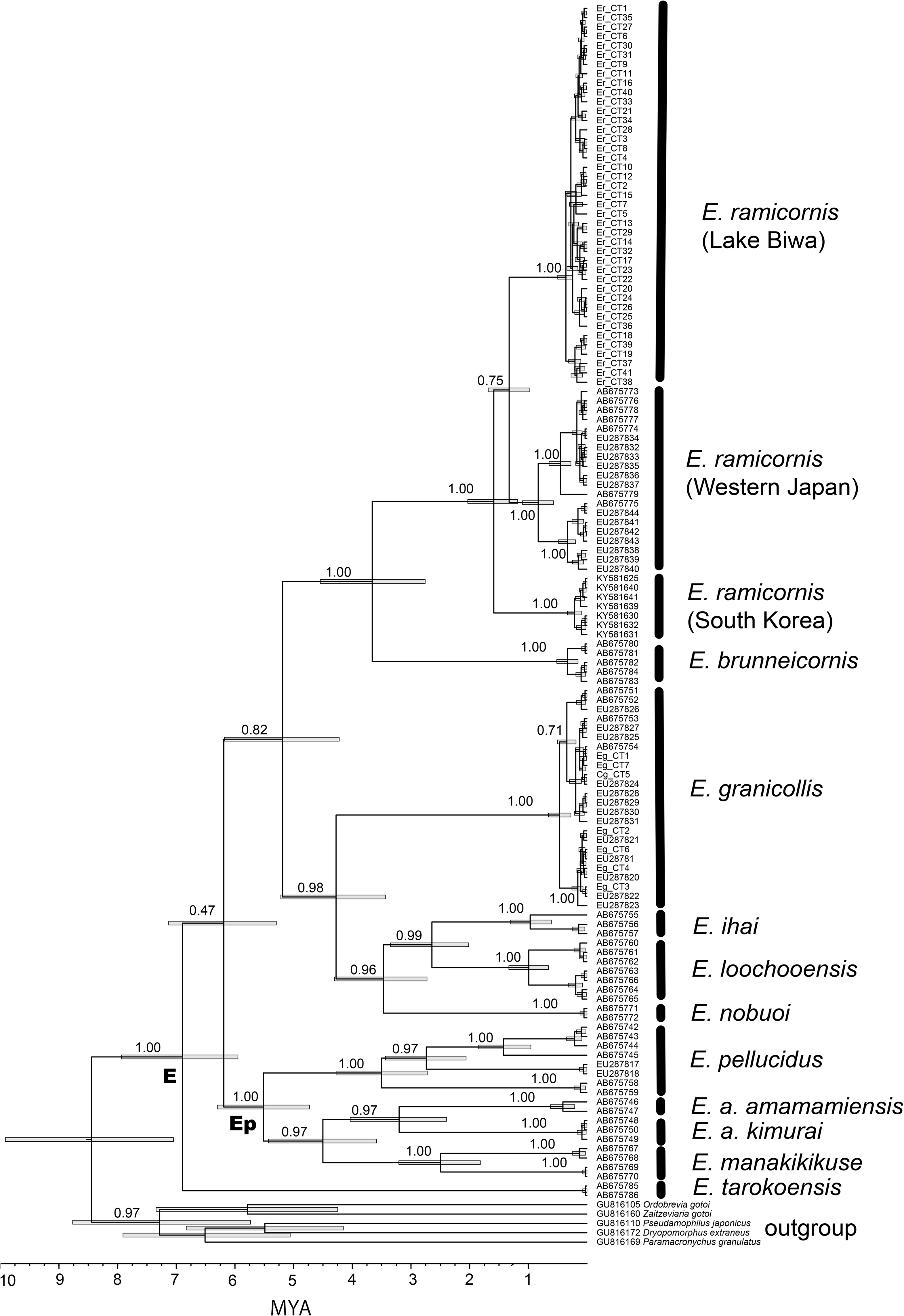
Divergence time of *Eubrianax* species inferred by Bayesian analysis and using 720-bp COI sequences. Topology is a tree with maximum reliability. Numbers at nodes indicate posterior probability. Probabilities for recent years are omitted. Calibration times used are indicated as “E” (*Eubrianax* group) and “Ep” (*E. pellucidus* species group) at nodes. Bars on nodes are 95% highest probability density (HPD) interval of node age. OTU names are shown as accession numbers or abbreviations of species name with haplotypes.

### Genetic distance

The phylogenetic analysis revealed that *E. ramicornis* displayed wide intraspecies divergence. To discuss the degree of differentiation, the genetic distance was calculated using 720-bp fragments of the COI “CT region” sequences within and between *E. ramicornis* groups and their related species *E. brunneicornis* and *E. granicollis*. The average genetic distances within the three *E. ramicornis* groups and their related species were in the range of 0.5–2.5% (Table 3). On the other hand, the pairwise genetic distances between the three species, *E. ramicornis*, *E. brunneicornis*, and *E. granicollis*, were in the range of 12.4–15.3%. The average genetic distances between the three *E. ramicornis* groups were in the range of 6.1–6.5% and they were intermediates values of those within groups/species and between three species.

**Table 3.** Average pairwise genetic distances within and between three groups of *E. ramicornis* and its relatives *E. brunneicornis* and *E. granicollis*. Abb.: abbreviations for *E. ramicornis* species or groups. Western Japan group does not include Lake Biwa.

|  | Group | Abb. | rLBw | rWJPN | rKOR | b | g |
| --- | --- | --- | --- | --- | --- | --- | --- |
| *E. ramicornis* | Lake Biwa | rLBw | 0.008 |  |  |  |  |
|  | western Japan | rWJPN | 0.061 | 0.025 |  |  |  |
|  | Korea | rKOR | 0.063 | 0.065 | 0.005 |  |  |
| *E. brunneicornis* |  | b | 0.130 | 0.132 | 0.124 | 0.009 |  |
| *E. granicollis* |  | g | 0.145 | 0.153 | 0.142 | 0.145 | 0.011 |

### Genetic differentiation within *E. ramicornis* of Lake Biwa

Sequence polymorphisms were detected within *E. ramicornis*. Therefore, we examined the sequence variations in detail. Here defined a haplotype as a unique sequence differing from others by ≥ 1 nucleotide. There were 30 LH region haplotypes in 81 individuals and 41 CT region haplotypes in 75 individuals (Figs. 3 and 4; Tables S3 and S4). There were more variations in the CT than the LH region of *E. ramicornis*. For the LH region, the frequency of the common LH2 type decreased from west to east along the lakeshore. Conversely, the common LH10 type was found mainly on the eastern to northern lakeshore. Network analysis of the LH haplotypes showed that the two most common haplotypes were LH2 and LH10, and the other minor haplotypes were derived from them (Fig. 5). LH2 and LH10 were found at collection sites from the western to the eastern lakeshore (Fig. 3). However, LH2 and all haplotypes derived from it except LH28 were found from the western to the northern shore whereas LH10 and all haplotypes derived from it were found from the northern to the eastern shore (Fig. 3 and 5).

**Figure 3.**
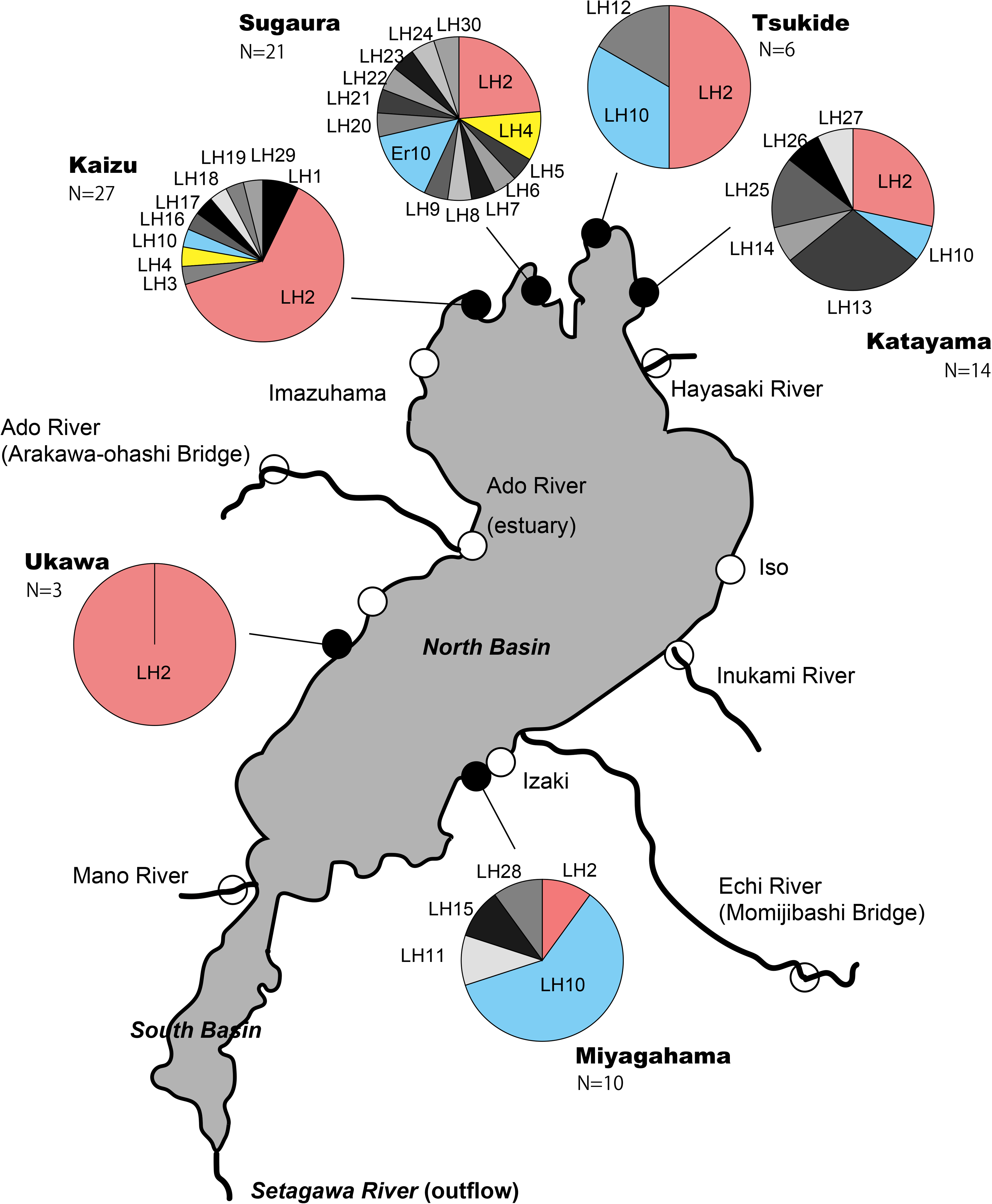
Distribution of *E. ramicornis* haplotypes found by using LH region of COI sequence. Bright colors show common haplotypes between sites. Grayscales show specific haplotypes at single sites. Circles on map indicate field sites. Black indicate sites where *E. ramicornis* founded. White indicates sites where *E. ramicornis* was absent.

**Figure 4.**
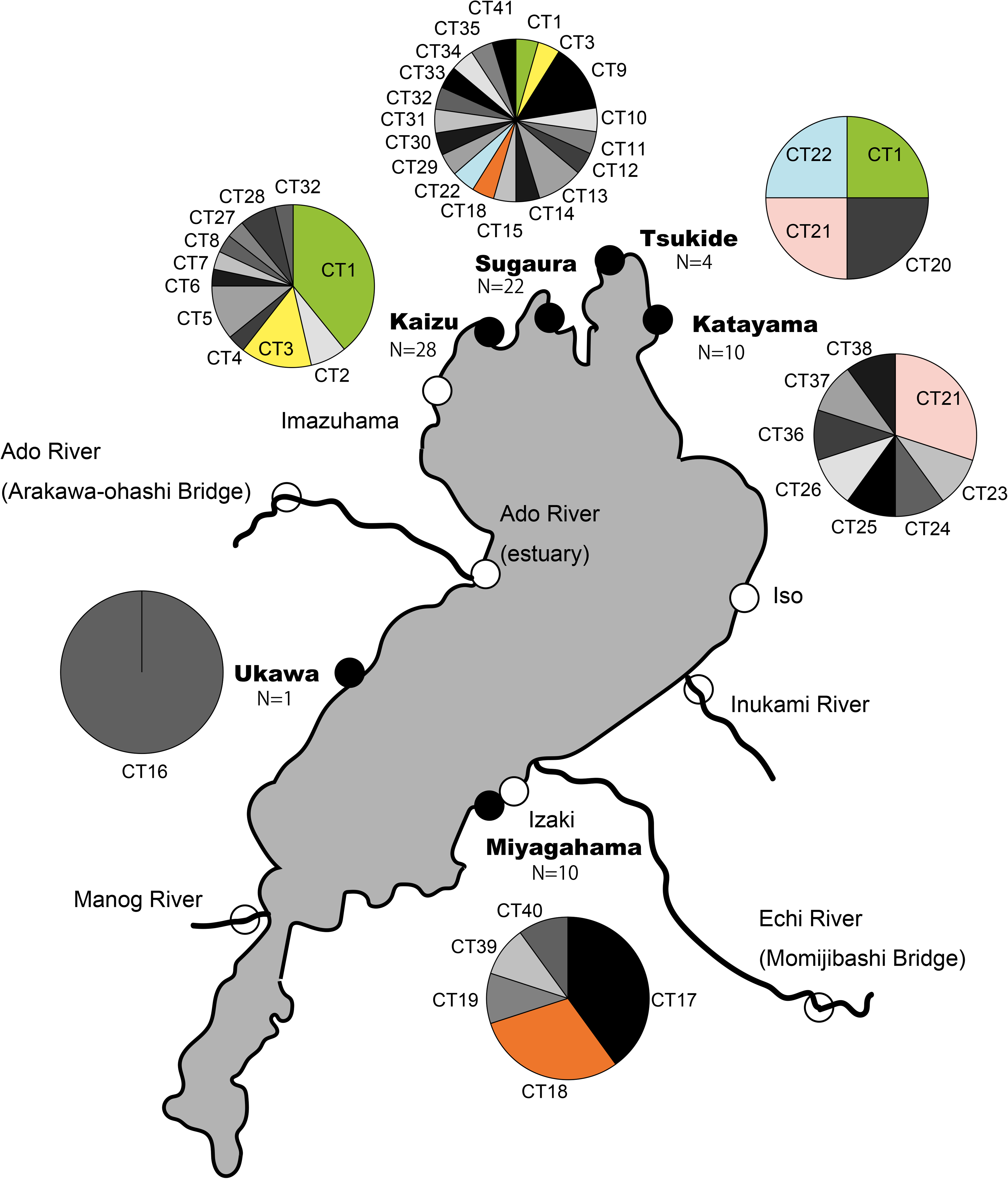
Distribution of haplotypes found in *E. ramicornis* using CT region of COI sequence. Descriptions of colors and sites are same as those for Fig. 3.

**Figure 5.**
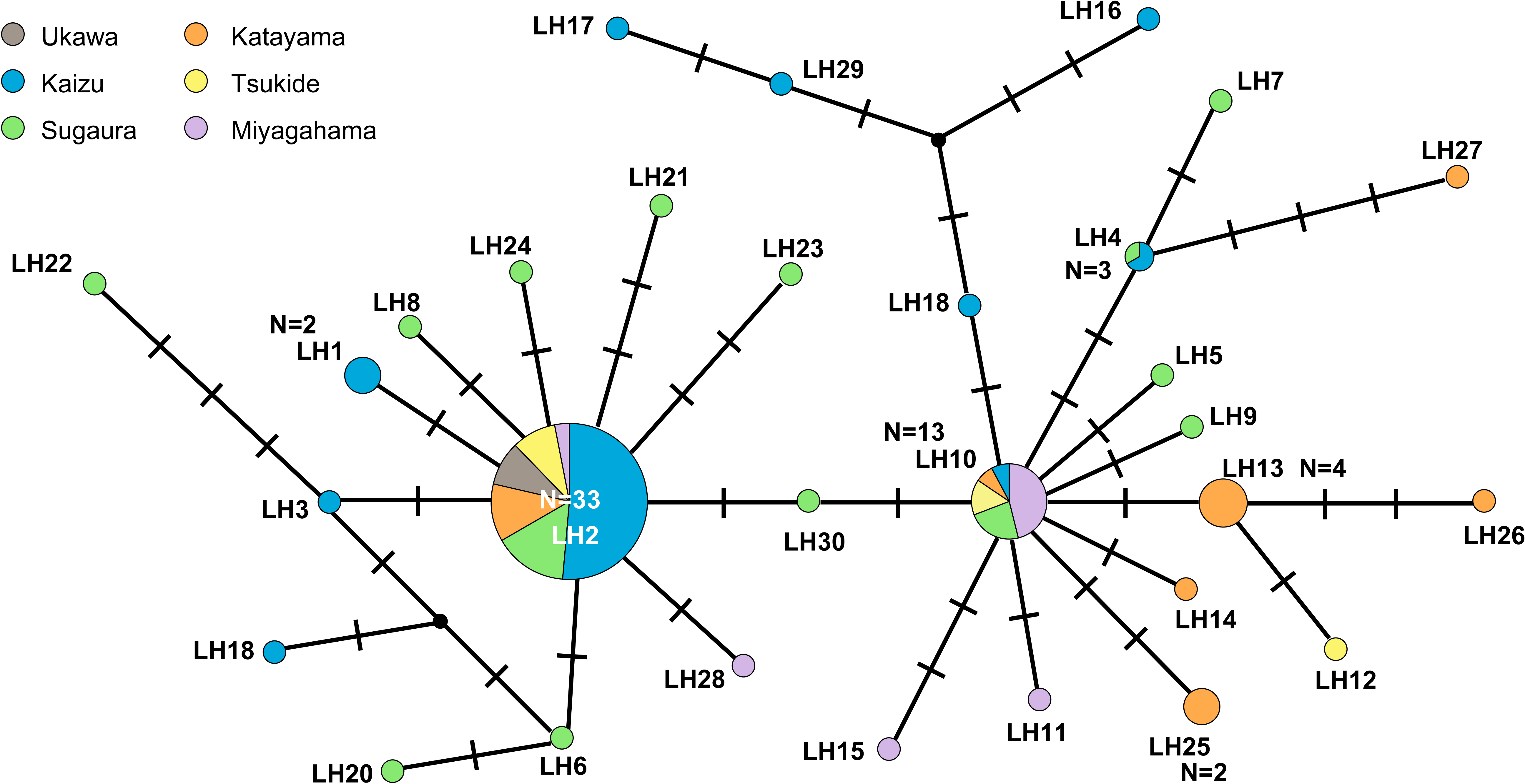
Haplotype network analyzed using LH region of COI in *E. ramicornis*. Colors indicate collection sites. Each circle represents one haplotype. Circle size increases with number of individuals. Number of each individual is one unless otherwise specified. Vertical lines represent single mutations. Black circles indicate assumed haplotypes.

The trends for the CT region were not as clear as those for the LH region as there were only a few individuals per haplotype (Fig. 4). However, all haplotypes except CT18 were relatively common to most adjacent collecting sites.

## Discussion

Psephenid larvae are generally found only in streams and small reservoirs. However, some have been detected in Lakes Biwa and Inawashiro in Japan and certain lakes in California (Murvosh, 1992; Nishino, 1992; Hayashi, 2009; Tsutsumi, 2017). The larvae of the four psephenid species collected in the present study usually live on and in gravelly and rocky fluvial materials (Hayashi, 2009). *E. granicollis* is found in large rivers whereas *E. ramicornis* is not. However, both species sympatrically occur in small-to medium-sized rivers (Hayashi, 2009). Here, *E. granicollis* was collected only from the rivers inflowing Lake Biwa while *E. ramicornis*, *Mal. japonicus*, and *Mat. japonicus* were collected from the gravelly and rocky shores of Lake Biwa (Table 2). These results were consistent with those of previous field studies on Lake Biwa and its watershed (Nishino, 1992; Hayashi, 2009; Japan Water Agency, 2018). Despite reports with uncertain species identification (Komatsu, 1964), both the present and previous studies suggested that habitats of *E. ramicornis* and *E. granicollis* may be separated by the shore and inflowing rivers of Lake Biwa, respectively. Three species excluding *E. ramicornis* were recorded also in the only outflowing river of Lake Biwa, namely, the Yodo River. Though *E. ramicornis* was not recorded there, four species including *E. ramicornis* were recorded in its tributaries (Hayashi, 2010; Tominaga, Shiyake & Beetle Team of Yodogawa River Research Group, 2012). In the present study, the sympatric occurrence of Psephenidae was observed only in Katayama where *E. ramicornis* and *Mal. japonicus* were collected. During the 1980’s, however, *E. ramicornis, Mal. japonicus*, and *Mat. japonicus* were sympatrically collected at > 10 sites on the lake shore in the North and South basins (Nishino, 1992). Several of our study sites overlapped with those reported in the 1980s. Nevertheless, we only collected *Mal. japonicus* and *Mat. japonicus* at Katayama and Iso, respectively (Table 2). Moreover, we detected no *Ectopria opaca* reported in previous studies (Hayashi, 2009; Japan Water Agency, 2018). Though this discrepancy may have been caused by sampling bias or insufficient sampling, there might nonetheless be a decline in psephenids along the shores of Lake Biwa. Despite the relatively lower present-day psephenid abundance, numerous individuals were collected from the northern shore of the North basin and many haplotypes were found there (Figs. 3 and 4). In the current North basin, the northern shore may be a more suitable habitat for psephenid larvae than the eastern or western shore. The water quality of the northern shore is conducive for psephenid larvae, the waves are high, and the rocks and gravel on the lake bottom are of appropriate size. These conditions create a comfortable environment for psephenid larvae. This tendency was consistent with the fact that the highest number of species of riverine insects was found on the northern shore of the North basin (Nishino et al., 2017).

*E. ramicornis* and *E. granicollis* have ecologically similar habitats and their geographical distributions overlap (Hayashi & Sota, 2008; Hayashi, 2009; Hayashi, 2010; Jung, Jach & Bae, 2020). However, only *E. ramicornis* has a genetically differentiated population in Lake Biwa (Fig. 1). Phylogenetic analyses revealed no genetic structures within the distribution ranges of *E. granicollis* and *E. ramicornis* in Japan except for Lake Biwa. Hence, gene flow covers a wide area (Fig. 1; Table S2). The average genetic distances within species were 2.5% for the *E. ramicornis* population of western Japan excluding Lake Biwa and 1.1% for *E. granicollis* (Table 3). The genetic distance within species was lower for *E. granicollis* than *E. ramicornis*. Thus, the former may have relatively higher migratory ability and/or recently expanded its distribution. *E. granicollis* might have, therefore, migrated into Lake Biwa more recently than *E. ramicornis*. This difference could explain the lack of genetic differentiation between the *E. granicollis* populations in the inflowing rivers of Lake Biwa and those elsewhere.

The average genetic distances between *E. ramicornis* populations in Lake Biwa (hereafter, “Lake Biwa group”), those in western Japan excluding Lake Biwa (hereafter, “western group”), and the Korean populations were all > 6% (Table 3). Both the Lake Biwa and western groups are located in Honshu Island, and the genetic distance between the groups is approximately equal to those between the Korean population across the sea. Though genetic distance is not a definitive species indicator, the apparent genetic differentiation shown by the Lake Biwa group suggests that it may be an independent species. Based on the currently available data, there are two possible hypotheses regarding the endemism of the Lake Biwa *E. ramicornis* group. It might either be endemic on its own or it belongs to a group that is genetically distinct from the western group. No definitive conclusion can be drawn from the results of the previous and present studies. On the other hand, the Lake Biwa group might be endemic based on its dispersal ability and geographical distance from other populations. The haplotypes at the six sites on the North basin were distributed along the shore from east to west via the northern shore (Figs. 3 and 4). A network analysis disclosed that the haplotypes were derived along the shore of Lake Biwa (Fig. 5). These results suggested that the Lake Biwa group has only limited dispersal ability. This group is more likely to move along the shore rather than disperse and migrate freely across the lake or towards the other watershed. Moreover, the geographical distances between the Lake Biwa group and neighboring populations are expected to be great because no *E. ramicornis* have been detected in the inflowing rivers of Lake Biwa in previous or present studies. The Lake Biwa group might be geographically isolated and unable to interact with other populations of *E. ramicornis*. Lake Biwa is surrounded by mountains and Lake Biwa group only occur in lakeshore but not the inflowing rivers or the outflowing river except for its tributaries. In order to test the above hypotheses, further analyses should be conducted with populations for which data are lacking in this study. These populations are those east of Lake Biwa and between Hyogo

Prefecture and Lake Biwa, which is the most eastern collection site of the western group treated in this study. Mating experiments between these populations would also provide further insight into the reproductive isolation factors that lead to the genetic differentiation obtained. No explicit morphological differences have so far been reported between the Lake Biwa and Western Japan groups.

The divergence of the Lake Biwa *E. ramicornis* group from the western group was estimated to have occurred ~1.78–1.07 MYA (median 1.40 MYA; Fig. 2) which corresponds to the Kusatsu Formation when Paleo-Lake Gamo disappeared (~1.8 MYA) followed by the appearance of Paleo-Lake Katata (~1.0 MYA) (Satoguchi, 2012; Satoguchi, 2020; Masuda & Satoguchi, 2021). It is believed that at that time, there were no lakes in the current location of the North basin of Lake Biwa and the entire area was repeatedly covered with swamps and flood plains. Geological research recently revealed that there used to be a lake on the eastern side of present-day Lake Biwa. It formed when the Yodo River tributary was dammed (Satoguchi, 2021). To the east, the Suzuka Mountains were raised during this period (Satoguchi, 2018). The area between the dammed lake and the Suzuka Mountains was Paleo-Lake Gamo. However, geological studies have shown that the disappearance of the fine mud-bottomed Paleo-Lake Gamo coincided with the differentiation of *E. ramicornis*. Furthermore, the ambient environment was fed gravel and sand from the Suzuka Mountains gradient (Satoguchi, 2018). Hence, this region would have been a suitable habitat for *E. ramicornis*. It is possible that *E. ramicornis* inhabited the gravel rivers that formed on the southeastern part of present-day Lake Biwa and was isolated from other local populations by the uplifted Suzuka Mountains and others surrounding Lake Biwa.

The timing of the evolution of the endemic animal species varied in Lake Biwa. Certain species differentiated during the Paleo-Lake period of the four-million-year history of Lake Biwa. However, others differentiated when Lake Biwa formed and deepened at its present location (Tabata et al., 2016; Saito et al., 2018; Hirano et al., 2019; Miura et al., 2019; Kaneko et al., 2021; Nishino, 2020). Divergence times estimated from fossil records and COI suggest that the fish species *Silurus lithophilus* and *Cottus reinii* (lacustrine type) endemic to Lake Biwa may have genetically differentiated in Lake Biwa during the same period as the Lake Biwa *E. ramicornis* group (Tabata et al., 2016). *Silurus lithophilus* inhabit northern rocky areas whereas other catfish species prefer areas with muddy bottoms and aquatic plants. *Cottus reinii* (lacustrine type) prefer the sand and gravel zones of the lakeshore and use the gravel bottoms of inflowing rivers for spawning (Tabata et al., 2016). The endemic mollusk *Valvata biwaensis* also diverged from its neighboring species 1.17–1.75 MYA (Saito et al., 2018). It preferentially inhabits sand-gravel over mud bottom. Other *Valvata* spp. are distributed across the rivers and lakes of northern Japan and are considered glacial relict species of northern origin. This finding is consistent with the fact that the Kusatsu Formation is regarded as a cold climate period. After Paleo-Lake Katata, present-day Lake Biwa emerged and expanded towards the present location. The climate warmed during that period but was interrupted by several glaciations. It is probable that the inflowing rivers had extremely low productivity, and related species immigrating from other areas could not establish there. These previous studies on other species that evolved during the same period suggested that geological changes such as the appearance of gravelly environments preferred by *E. ramicornis*, geological isolation caused by orogenic activity and mountain uplift, and dispersal limitation caused by climate changes contributed to the mechanism of the genetic differentiation of the Lake Biwa *E. ramicornis* group.

Little is known about the mechanism of genetic differentiation of aquatic insects in Lake Biwa. Only the caddisfly *A. biwaensis* and the mayfly *E. limnobium* endemic to Lake Biwa have been described thus far (Nishimoto, 1994; Ishiwata, 1996). Nevertheless, the times at which they diverged from related species are unknown. The white mayfly *E. yoshidae* is not endemic to Lake Biwa but the time of divergence of the Lake Biwa clade from its nearest relative in Tohoku was estimated to be ~1.0 MYA (Kaneko et al., 2021).

There may be other examples of aquatic insect species with a wide distribution that are genetically differentiated in Lake Biwa. Examples of genetic differentiation in Lake Biwa have also been reported for other widely distributed species other than aquatic insects. The ayu fish *Plecoglossus altivelis* subsp. and the lake prawn *Palaemon paucidens* are widely distributed across Japan and in Lake Biwa. However, the populations at Lake Biwa ecologically and genetically differ from other local populations (Azuma, 1973; Nishino, 1980; Nishida, 1985, 1986; Chow, Fujino & Nomura 1988; Chow et al., 2018). The high relative mobility of the aquatic insect populations in Lake Biwa and possibly the lack of sufficient studies on them may explain why it difficult to differentiate them from those in other areas. Phylogenetic analysis of common, widely distributed aquatic insect species may help elucidate the mechanisms by which they genetically differentiated at Lake Biwa.

The Lake Biwa *E. ramicornis* group inhabits a lake which is a lentic environment. This population has low mobility and was genetically differentiated from those of other populations. These results apparently contradict the habitat stability hypothesis which states that organisms inhabiting lentic waters have high dispersal capacity and low genetic structure (Ribera & Vogler, 2000; Ribera, 2008). Several studies revealed that barriers to migration and habitat stability in long-term, historical, and geological backgrounds and rather than water flow strongly affects genetic differentiation in aquatic insects (Letsch, Gottsberger & Ware, 2016; Saito & Tojo, 2016; Tojo et al., 2017; Takenaka et al., 2021). Genetic differentiation of the Lake Biwa *E. ramicornis* group could also be explained without the habitat stability hypothesis. The Lake Biwa *E. ramicornis* group might have genetically differentiated from its ancestral population because of the geographic history of Lake Biwa formation. Moreover, the lakeshore inhabited by *E. ramicornis* is not truly lentic as it is widely continuous and stable. Rocky and sandy-gravelly lakeshores accounts for 17% and ~30%, respectively, of the total 235 km length of Lake Biwa (Tatsumi, 2017). Nishino (2017) classified Lake Biwa into two main regions based on their geographic characteristics and organism distribution, namely, “Lake Biwa as a great lake” and “Lake Biwa as a swamp”. The former is sandy, steeply sloping rocky lakeshore with prevailing wind and waves where numerous endemic species occur. This environment is the preferred habitat of *E. ramicornis* and its water kinesis resembles that of a river rather than that of an unstable, motionless lentic water body.

In this study, all the specimen and sequence data other than Lake Biwa were obtained from previously published research (Hayashi & Sota, 2008; Hayashi, Song & Sota, 2012; Jung, Jach & Bae, 2020) and GenBank. This limited the analysis of phylogenetic analysis and genetic distance, as only the CT region of the COI 3’-end of the mitochondrial gene in the database could be used. As already shown by Hayashi, Song & Sota (2012), *E. ramicornis* collected west of Lake Biwa was clearly divided into two lineages, but there was no geographical tendency between these two lineages (Fig. 1). Hayashi, Song & Sota (2012) did not specifically mention this branching in the text, other than in the figure. One possibility for this branching is that biases in population dynamics due to cytoplasmic factors, such as infection by intracellular symbiotic bacteria such as Wolbachia, may influence genetic differentiation (Werren, Baldo & Clark, 2008). Analysis of the nuclear genes of *E. ramicornis* and investigation of the bacterial infections in this species may disclose certain factors accounting for the genetic differences between the *E. ramicornis* populations inside and outside Lake Biwa. The latter remains the standard and most popular region used in DNA barcoding (Hebert et al., 2003a; Hebert, Ratnasingham & de Waard, 2003b). On the other hand, the LH region, which was only analysed in the Lake Biwa Group in this study, is the standard and most common region used for DNA barcoding (Hebert et al., 2003a; Hebert, Ratnasingham & de Waard, 2003b). DNA barcodes are sequences of specific gene regions suitable for identifying species and are widely used to identify morphologically difficult-to-identify species and to detect biodiversity using environmental DNA (Hajibabaei et al., 2019; Uchida et al., 2020; Ficetola et al., 2021). In particular, conventional collection surveys of aquatic insects require a great deal of time and expertise in identification. Environmental DNA-based surveys have attracted attention as a way of compensating for this. An enhanced database of DNA barcodes for referencing the sequences obtained is essential to improve the sensitivity and accuracy of the results of environmental DNA-based diversity surveys. However, to date, only a few species have been registered in the LH region of the Psephenids, and this is the first registration for the four species obtained in this study (as of November 14^th^, 2022). We expect that the LH region sequences obtained in this study will be widely applied in aquatic biodiversity monitoring and research.

The 2015 edition of the Shiga Prefecture Red Data Book listed 56% of the endemic species in Lake Biwa as endangered or rare (Nishino, 2018). In considering biological conservation measures in Lake Biwa, it is important to understand the endemicity and evolutionary process of each organism. However, only two endemic species of aquatic insects are known, and there are few examples of studies of genetic relationships between Lake Biwa and outside of Lake Biwa (Kaneko et al., 2021). It is important to urgently figure out if there are hidden endemic species in aquatic insects as well, and if necessary, consider conservation measures. Lake Biwa is one of only about 20 ancient lakes in the world. In other ancient lakes, many endemic species have been reported for fish, mollusks, and amphipods, but knowledge of endemic aquatic insects is as limited as in Lake Biwa. Tabata et al. (2016) proposed that the evolutionary path of the paleo-Lake Biwa transition during the formation of present Lake Biwa was different from the events that occurred in other typical ancient lakes, and therefore the evolutionary course of organisms may be different as well. The examples of *E. ramicornis* and other aquatic insects in Lake Biwa shown in this study may suggest that other ancient lakes also hide endemic or genetically differentiated aquatic insect species that are simply not highlighted. Or it may suggest a unique phenomenon seen in Lake Biwa, which has repeatedly disappeared, emerged, and migrated over a period of 4 million years. We expect that this study will contribute to the study of the evolution of aquatic insects in ancient lakes and to conservation measures.

## Conclusion

The present study confirmed that the Lake Biwa *E. ramicornis* population was genetically differentiated from other *E. ramicornis* populations in western Japan. However, no morphological differences between these populations were detected. Nevertheless, the estimated genetic distances and divergence times were valid and could suffice to indicate speciation. When the Lake Biwa *E. ramicornis* population genetically diverged, the region was characterized by recurring lakes and marshes/floodplains and was, therefore, unstable. On the other hand, geological studies suggested that the region included a gravelly riverine environment that *E. ramicornis* originally preferred as its habitat. Geographic barriers such as the surrounding faults and provenances and the newly uplifted mountain ranges might account for the genetic differentiation between the Lake Biwa *E. ramicornis* population and the others. Although the closely related species *E. granicollis* prefers habitats similar to those inhabited by *E. ramicornis*, its populations at Lake Biwa and other areas were not genetically differentiated. This discrepancy may be explained by their habitat preferences and the timing of their invasion in the Lake Biwa basin. Future nuclear gene and genome-wide polymorphism analyses will validate this hypothesis. Genetic analyses may reveal hidden genetic differentiation and widely distributed endemic species across Lake Biwa and other regions. There is also little knowledge on the genetic endemism of aquatic insect species in other ancient lakes. Future research is awaited to determine whether the genetic differentiation observed in Lake Biwa is a common phenomenon in other ancient lakes.

## Acknowledgements

We thank Y. Sakai and S. Ishiwata for their help of sampling, J. Machizawa, T. Shindo and M.
Nakajima for their technical assistance. This work was supported by the Collaborative Research Fund from Shiga Prefecture “Studies on conservation and ecosystem management of Lake Biwa” under the Japanese Grant for Regional Revitalization and JSPS KAKENHI Grant Number 21H03624.

## SUPPLEMENTARY INFORMATION

**Table S1. Annealing temperature, PCR primers, and number of PCR cycles used in the present study.**

**Table S2. COI sequences of the CT region used in the phylogenetic analyses of the present study.**

**Table S3. Psephenid species collected in the present study and their haplotypes or accession numbers for two COI regions.** Haplotypes are shown for *E. granicollis* and *E. ramicornis*. Accession numbers are shown for other species and Bw199. No haplotype name was assigned for LH of Bw199 or CT of BW197 as both sequences were shorter than all others. See Table S4 for haplotype accession number. nd: not detected.

**Table S4. Haplotypes and accession numbers of the two COI regions.**

## References

Abellan P, Millan A, Ribera I. 2009. Parallel habitat-driven differences in the phylogeographical structure of two independent lineages of Mediterranean saline water beetles. Molecular Ecology 18: 3885–3902 DOI: 10.1111/j.1365-294X.2009.04319.x.

Arribas P, Velasco J, Abellan P, Sanchez-Ferrnandez D, Andujar C, Calosi P, Millan A, Ribera I, Bilton D. 2012. Dispersal ability rather than ecological tolerance drives differences in range size between lentic and lotic water beetles (Coleoptera: Hydrophilidae). Journal of Biogeography 39: 984–994 DOI: 10.1111/j.1365-2699.2011.02641.x.

Azuma M. 1973. Studies on the variability of the landlocked ayu-fish, *Plecoglossus altivelis* T. et S., in Lake Biwa IV. Considerations on the grouping and features of viability. Japanese Journal of Ecology 23: 255–265 DOI: 10.18960/seitai.23.4_147.

Bandelt H-J, Forster P, Sykes BC, Richards MB. 1995. Mitochondrial portraits of human populations. Genetics 141:743–753 DOI: 10.1093/genetics/141.2.743.

Bouckaert R, Vaughan TG, Barido-Sottani J, Duchêne S, Fourment M, Gavryushkina A, Heled J, Jones G, Kühnert D, De Maio N, Matschiner M, Mendes FK, Müller NF, Ogilvie HA, du Plessis L, Popinga A, Rambaut A, Rasmussen D, Siveroni I, Suchard MA, Wu CH, Xie D, Zhang C, Stadler T, Drummond AJ. 2019. BEAST 2.5: An advanced software platform for Bayesian evolutionary analysis. PLoS Computational Biology 15: e1006650 DOI: 10.1371/journal.pcbi.1006650.

Budzakoska-Gjoreska B, Trajanovski S, Trajanovska S. 2014. Comparative biocenological analysis of Gastropoda on the Macedonian part of Lake Ohrid and its watershed. Biologia 69: 1023–1029 DOI: 10.2478/s11756-014-0403-7.

Chow S, Fujino Y, Nomura T. 1988. Reproductive isolation and distinct population structures in two types of the freshwater shrimp *Palaemon paucidens*. Evolution 42: 804–813 DOI: 10.1111/j.1558-5646.1988.tb02498.x.

Chow S, Imai T, Ikeda M, Maki S, Ookuni T, Muto F, Nohara K, Furusawa C, Shichiri H, Nigorikawa N, Uragaki N, Kawamura A, Ichikawa T, Ushioda K, Higuchi M, Tega T, Kodama K, Itoh M, Ichimura M, Matsuzaki K, Hirasawa K, Tokura K, Nakahata K, Kosama S, Hakoyama H, Yada T, Niwa K, Nagai S, Yanagimoto T, Saito K, Nakaya M, Maruyama T. 2018. A DNA marker to discriminate two types of freshwater shrimp *Palaemon paucidens* and the distribution of these two types in Japan. Nippon Suisan Gakkaishi 84: 674–681 DOI: 10.2331/suisan.17-00082. (in Japanese with English summary).

Cristescu ME, Adamowicz SJ, Vaillant JJ, Haffner DG. 2010. Ancient lakes revisited: from the ecology to the genetics of speciation. Molecular Ecology 19: 4837–4851. DOI: 10.1111/j.1365-294X.2010.04832.x.

Dijkstra K-DB, Monaghan MT, Pauls SU. 2014. Freshwater biodiversity and aquatic insect diversification. Annual Reviews of Entomology 59: 143–63. DOI: 10.1146/annurev-ento-011613-161958.

Drummond R and Bouckaert RR. 2015. Bayesian Evolutionary Analysis with Beast. Cambridge University Press.

Ficetola GF, Boyer F, Valentini A, Bonin A, Meyer A, Dejean T, Gaboriaud C, Usseglio-Polatera P, Taberlet P. 2021. Comparison of markers for the monitoring of freshwater benthic biodiversity through DNA metabarcoding. Molecular Ecology 30: 3189–3202 DOI: 10.1111/mec.15632.

Fluxus Engineering. 2020. Network 10.1.0.0 Available at https://www.fluxus-engineering.com/sharenet.htm (accessed 27 July 2020).

Folmer O, Black M, Hoeh W, Lutz R, Vrijenhoek R 1994. DNA primers for amplification of mitochondrial cytochrome *c* oxidase subunit I from diverse metazoan invertebrates. Molecular Marine Biology and Biotechnology 3: 294–299.

Grabowski M, Wysocka A, Mamos T. 2017. Molecular species delimitation methods provide new insight into taxonomy of the endemic gammarid species flock from the ancient Lake Ohrid. Zoological Journal of the Linnean Society 181: 272–285 DOI: 10.1093/zoolinnean/zlw025.

Gullan PJ, Cranston PS. 1994 The Insects: an outline of entomology. London, UK: Chapman and Hall, 230–233.

Gurkov A, Rivarola-Duarte L, Bedulina D, Casas IF, Michael H, Drozdova P, Nazarova A, Govorukhina E, Timofeyev M, Stadler PF, Luckenbach T. 2019. Indication of ongoing amphipod speciation in Lake Baikal by genetic structures within endemic species. BMC Evolutionally Biology 19: 138 DOI: 10.1186/s12862-019-1470-8.

Hajibabaei M, Porter TM, Robinson CV, Baird DJ, Shokralla S, Wright MTG. 2019. Watered-down biodiversity? A comparison of metabarcoding results from DNA extracted from matched water and bulk tissue biomonitoring samples. Plos One 14: e0225409 DOI: 10.1371/journal.pone.0225409.

Hayashi M. 2009. Studies on the Japanese members of Psephenidae. Bulletin of the Hoshizaki Green Foundation 12: 35–85. (in Japanese)

Hayashi M. 2010. Distributional records of Psephenidae in Japan. Bulletin of the Hoshizaki Green Foundation 13: 301–322. (in Japanese)

Hayashi M, Kawakami Y. 2009. Fossil of the genus *Eubrianax* (Coleoptera, Psephenidae) from the Upper Miocene Ningyotoge Formation in Tottori Prefecture, Japan. Elytra 37: 99–103.

Hayashi M. and Sota T. 2008. Discrimination of two Japanese water pennies, *Eubrianax granicollis* Lewis and *E. ramicornis* Kisenwetter (Coleoptera: Psephenidae)), based on laboratory rearing and molecular taxonomy. Entomological Science 11: 349–357 DOI: 10.1111/j.1479-8298.2008.00277.x.

Hayashi M, Song SD, Sota T. 2012. Molecular phylogeny and divergence of the Water penny genus *Eubrianax* (Coleoptera: Psephenidae) in Japan. Entomological Science 15: 314–323 DOI: 10.1111/j.1479-8298.2012.00518.x.

Hebert PDN, Cywinska A, Ball SL, deWaard JR. 2003a. Biological identifications through DNA barcodes. Proceedings of the Royal Society of London. Series B: Biological Sciences 270: 313–321 DOI: 10.1098/rspb.2002.2218.

Hebert PDN, Ratnasingham S, deWaard JR. 2003b. Barcoding animal life: cytochrome *c* oxidase subunit 1 divergences among closely related species. Proceedings of the Royal Society of London. Series B: Biological Sciences 270: S96–S99. DOI: 10.1098/rsbl.2003.0025.

Hirano T, Saito T, Tsunamoto Y, Koseki J, Prozorova L, Do VT, Matsuoka K, Nakai K, Suyama Y, Chiba S. 2019. Role of ancient lakes in genetic and phenotypic diversification of freshwater snails. Molecular Ecology 28: 5032–5051 DOI: 10.1111/mec.15272.

Ichise S, Sakamaki Y, Shimano SD. 2021. Neotypification of *Difflugia biwae* (Amoebozoa: Tubulinea: Arcellinida) from the Lake Biwa, Japan. Species Diversity 26: 171–186 DOI: 10.12782/specdiv.26.171.

Ishiwata S. 1996. A study of the genus *Ephoron* from Japan (Ephemeroptera, Polymitarcyidae). Canadian Entomologist 128: 551–572.

Ishiwata S, Uenishi M. 2020. Distribution of *Ephoron shigae* (Takahashi, 1924) in Shiga Prefecture. Biology of Inland Waters 35: 81–84 (in Japanese)

Japan Water Agency. 2018. The investigation report of benthic animals in Lake Biwa with pictures, 2nd ed. https://www.water.go.jp/kansai/biwako/html/download/files/zusetsu/2018033002_benthos.pdf (accessed 21 March 2022) (in Japanese)

Jung SW, Jach MA, Bae YJ. 2020. Review of the Water penny beetles (Coleoptera: Psephenidae) of the Korean Peninsula based on morphology and mitochondrial cytochrome *c* oxidase subunit I gene sequences. Journal of Asia-Pacific Biodiversity 13: 13–23 DOI: 10.1016/j.japb.2019.10.003.

Kaneko H, Ishiwata S, Bae YJ, Takamura-Enya T. 2021. Genetic characteristics and phylogeography of the habitat generalist mayfly *Ecdyonurus yoshidae* (Ephemeroptera: Heptageniidae) in the Japanese archipelago. Entomological Research 51: 238–250 DOI: 10.1111/1748-5967.12498.

Kawabe, T. 1994. Biwako no oitachi (Formation of Lake Biwa). In: Research Group for Natural History of Lake Biwa ed. Biwako no shizenshi (The natural history of Lake Biwa). Tokyo: Yasaka Shobo, 24–72. (in Japanese)

Komatsu T. 1964. Aquatic insect communities in winter and the biotic index of the rivers which flow into the Lake Biwa. Japanese Journal of Ecology 14: 217–223.

Kontula T, Kirilchik SV, Väinölä R. 2003. Endemic diversification of the monophyletic cottoid fish species flock in Lake Baikal explored with mtDNA sequencing. Molecular Phylogenetics and Evolution 27: 143–155 DOI: 10.1016/S1055-7903(02)00376-7.

Koroiva R, Pepinelli M. 2019. Distribution and Habitats of Aquatic Insects. In: Del-Claro K & Guillermo R eds Aquatic Insects: Behavior and Ecology. Cham: Springer. DOI: 10.1007/978-3-030-16327-3_2.

Kumar S, Stecher G, Li M, Knyaz C, Tamura K. 2018. MEGA X: Molecular Evolutionary Genetics Analysis across computing platforms. Molecular Biology and Evolution 35: 1547–1549 DOI: 10.1093/molbev/msy096.

Lee CF, Yang PS, Sato M. 2001. Phylogeny of the genera of Eubrianaciae and description of additional members of *Eubrianax* (Coleoptera: Psephenidae). Annales of the Entomological Society of America 94: 347–362.

Letsch H, Gottsberger B, Ware JL. 2016. Not going with the flow: a comprehensive time-calibrated phylogeny of dragonflies (Anisoptera: Odonata: Insecta) provides evidence for the role of lentic habitats on diversification. Molecular Ecology 25: 1340–1353 DOI: 10.1111/mec.13562.

Maehata M. 2020. Characteristics of the Ichthyotauna of Lake Biwa, with special reference to its long-term changes. In Kawanabe, H., M. Nishino and M. Maehata eds. Lake Biwa: Interactions between Nature and People 2^nd^ ed. Cham: Springer.

Marten A, Brandle M, Brandle R. 2006. Habitat type predicts genetic population differentiation in freshwater invertebrates. Molecular Ecology 15: 2643–2651 DOI: 10.1111/j.1365-294X.2006.02940.x.

Martens K. 1997. Speciation in ancient lakes. Trends in Ecology and Evolution 12: 177–182 DOI: 10.1016/s0169-5347(97)01039-2.

Masuda F, Satoguchi Y. 2021. New findings concerning the paleoenvironmental change of Lake Biwa based on the results of facies analysis for the Karasuma Deep Core. Research Report of the Lake Biwa Museum 34: 95–109. DOI: 10.51038/rrlbm.34.0_95. (in Japanese)

Matsuoka K. 1987. Malacofaunal succession in Pliocene to Pleistocene non-marine sediments in the Omi and Ueno Basins, Central Japan. Journal of Earth Sciences 35: 23–115.

Miura O, Urabe M, Nishimura T, Nakai K, Chiba S. 2019. Recent lake expansion triggered the adaptive radiation of freshwater snails in the ancient Lake Biwa. Evolution Letters 3: 43–54. DOI: 10.1002/evl3.92.

Murvosh CM. 1992. On the occurrence of the Water penny beetle *Eubrianax edwardsii* in lentic ecosystems (Coleoptera: Psephenidae). The Coleopterists Bulletin 46: 43–51.

Nishida M. 1985. Substantial genetic differentiation in ayu *Plecoglossus altivelis* of the Japan and Ryukyu Islands. Bulletin of the Japanese Society of Scientific Fisheries 51:1269–1274 DOI: 10.2331/suisan.51.1269.

Nishida M. 1986. Geographic variation in the molecular, morphological and reproductive characters of the ayu *Plecoglossus altivelis* (Plecoglossidae) in the Japan-Ryukyu Archipelago. Japanese Journal of Ichthyology 33: 232–248 DOI: 10.11369/jji1950.33.232.

Nishimoto H. 1994. A new species of *Apatania* (Trichoptera, Limnephilidae) from lake Biwa, with notes on its morphological variation within the lake. Japanese Journal of Entomology 62: 775–785.

Nishino M. 1980. Geographical variations in body size, brood size and egg size of a freshwater shrimp, *Palaemon paucidens* de Haan, with some discussion on brood habit. Japanese Journal of Limnology 41: 185–202.

Nishino M. 1992. Biwako no teisei-dobutsu: Mizube no ikimono-tachi. II. Suisei-konchu hen. (Benthic Animals in Lake Biwa: animals of the Water Shore. II Aquatic Insects) Otsu: Lake Biwa Research Institute, 52–54. (in Japanese)

Nishino M., Akiyama M, Nakajima T. 2017. Messages from Lake Biwa Shore: viewpoints of its conservation and restoration. Hikone: Sunrise Publishing. (in Japanese)

Nishino M. 2018 Koyushu (Endemic species of Lake Biwa). In Shiga Prefectural Government eds. Biwako Handbook 3^rd^ ed. (Handbook of Lake Biwa). pp 152–153. (in Japanese)

Nishino M. 2020. Biodiversity of Lake Biwa and Adjacent Areas. In Kawanabe H, Nishino M, Maehata M. eds. Lake Biwa: Interactions between Nature and People 2^nd^ ed. Cham: Springer, 71–81.

Polzin T, Daneschmand SV. 2003. On Steiner trees and minimum spanning trees in hypergraphs. Operations Research Letters 31:12–20 DOI: 10.1016/S0167-6377(02)00185-2.

Rambaut A. 2012. FigTree Tree Figure Drawing Tool Version 1.4.3. Available at http://tree.bio.ed.ac.uk/software/figtree/

Rambaut A, Drummond AJ, Xie W, Baele G, Suchard MA. 2018. Tracer MCMC Trace Analysis Tool Version 1.7.1. Available at http://tree.bio.ed.ac.uk/softwar/tracer/

Ribera I. 2008. Habitat constraints and the generation of diversity in freshwater macroinvertebrates. Lancaster J, Briers RA. eds. In Aquatic Insects: Challenges to Populations: Proceedings of the Royal Entomological Society’s 24th Symposium. Royal Entomological Society of London. Symposium. Oxford: CAB International.

Ribera I, Vogler AP. 2000. Habitat type as a determinant of species range sizes: The example of lotic-lentic differences in aquatic Coleoptera. Biologoical Journal of the Linnean Society 71: 33–52 DOI: 10.1006/bijl.1999.0412.

Ribera I, Bilton DT, Vogler AP. 2003. Mitochondrial DNA phylogeography and population history of Meladema diving beetles on the Atlantic Islands and in the Mediterranean basin (Coleoptera, Dytiscidae). Molecular Ecology 12: 153–167 DOI: 10.1046/j.1365-294x.2003.01711.x.

Saito T, Prozorova L, Sitnikova T, Surenkhorloo, Hirano T, Morii Y, Chiba S. 2018. Molecular phylogeny of glacial relict species: a case of freshwater Valvatidae molluscs (Mollusca: Gastropoda) in North and East Asia. Hydrobiologia 818: 105–118 DOI: 10.1007/s10750-018-3595-y.

Saito R, Tojo K. 2016. Comparing spatial patterns of population density, biomass, and genetic diversity patterns of the habitat generalist mayfly *Isonychia japonica* Ulmer (Ephemeroptera: Isonychiidae) in the Chikuma-shinano River basin. Freshwater Science 35: 712–723.

Satoguchi Y. 2012. Geological history of Paleo- and present Lake Biwa. In: Kawanabe H, Nishino M., Maehata M. eds. Lake Biwa: Interactions between Nature and People. Cham: Springer, 9–16.

Satoguchi Y. 2018. When was Lake Biwa formed? Shiga: Sunrise Publishing Co., Ltd. (in Japanese)

Satoguchi Y. 2020. Geological history of paleo- and present Lake Biwa. In Kawanabe H, Nishino M., Maehata M. eds. Lake Biwa: Interactions between Nature and People 2^nd^ ed. Cham: Springer, 17–24, Springer.

Sawada N, Nakano T. 2021. Revisiting a 135-year-old taxonomic account of the freshwater snail *Semisulcospira multigranosa:* designating its lectotype and describing a new species of the genus (Mollusca: Gastropoda: Semisulcospiridae). Zoological Studies 60:7 DOI: 10.6620/ZS.2021.60-07.

Seehausen O. 2002. Patterns in fish radiation are compatible with Pleistocene desiccation of Lake Victoria and 14 600 year history for its cichlid species flock. Proceedings. Biological sciences / The Royal Society 269: 491–497 DOI: 10.1098/rspb.2001.1906.

Seehausen O. 2006. African cichlid fish: a model system in adaptive radiation research *Proceedings*. Biological sciences / The Royal Society 273: 1987–1998 DOI: 10.1098/rspb.2006.3539.

Sekiné K, Hayashi F, Tojo K. 2013. Phylogeography of the East Asian polymitarcyid mayfly genus *Ephoron* (Ephemeroptera: Polymitarcyidae): a comparative analysis of molecular and ecological characteristics. Biological Journal of the Linnean Society 109: 181–202 DOI: 10.1111/bij.12033.

Simon C, Frati F, Beckenbach A, Crespi B, Liu H, Flook P. 1994. Evolution, weighting, and phylogenetic utility of mitochondrial gene sequences and a compilation of conserved polymerase chain reaction primers. Annals of the Entomological Society of America 87: 651–701 DOI: 10.1093/aesa/87.6.651.

Sota T, Hayashi M. 2007. Comparative historical biogeography of *Plateumaris* leaf beetles (Coleoptera: Chrysomelidae) in Japan: intberplay between fossil and molecular data. Journal of Biogeography 34: 977–993 DOI: 10.1111/j.1365-2699.2006.01672.x.

Stelbrink B, Shirokaya AA, Föller K, Wilke T, Albrecht C. 2016. Origin and diversification of Lake Ohrid’s endemic acroloxid limpets: the role of geography and ecology. BMC Evolutionary Biology 16: 273 DOI: 10.1186/s12862-016-0826-6.

Tabata R, Kakioka R, Tominaga K, Komiya T, Watanabe K. 2016. Phylogeny and historical demography of endemic fishes in Lake Biwa: the ancient lake as a promoter of evolution and diversification of freshwater fishes in western Japan. Ecology and Evolution 6: 2601–2623 DOI: 10.1002/ece3.2070.

Takenaka M, Shibata S, Ito T, Shimura N, Tojo K. 2021. Phylogeography of the northernmost distributed *Anisocentropus* caddisflies and their comparative genetic structures based on habitat preferences. Ecology and Evolution 11: 4957–4971 DOI: 10.1002/ece3.7419.

Tatsumi M. 2017. Characteristics and current status of the lakeshore landscapes of Lake Biwa. In Nishino M. ed Message from the shore of Lake Biwa - Perspectives on conservation and restoration -. Shiga: Sunrise Publishing Co., Ltd., 50–62. (in Japanese)

Tojo K, Sekine K, Takenaka M, Isaka Y, Komaki S, Suzuki T, Schoville SD. 2017. Species diversity of insects in Japan: Their origins and diversification processes. Entomological Science 20: 357–380 DOI: 10.1111/ens.12261.

Tominaga O, Shiyake S, Beetle Team of Yodogawa River Research (Project Y2, Osaka Museum of Natural History). 2012. Fauna and distribution of the family Psephenidae in the Yodogawa River system, central Japan. Bulletin of the Osaka Museum of Natural History, 66: 39–48. (in Japanese)

Tsutsumi T. 2017. Preliminary notes on the macrobenthic invertebrate fauna of Lake Inawashiro, Fukushima Prefecture, Japan. Jorunal of Center for Regional Affairs, Fukushima University 28: 57–71.

Uchida N, Kubota K, Aita S, Kazama S. 2020. Aquatic insect community structure revealed by eDNA metabarcoding derives indices for environmental assessment. PeerJ 8: e9176 DOI: 10.7717/peerj.9176.

Verheyen E, Walter S, Snoeks J, Axel M. 2003. Origin of the Superflock of Cichlid Fishes from Lake Victoria, East Africa. Science 300: 325–329 DOI: 10.1126/science.1080699.

Werren J, Baldo L, Clark M. 2008. *Wolbachia:* master manipulators of invertebrate biology. Nature Reviews. Microbiology 6: 741–751 DOI: 10.1038/nrmicro1969.

