## Supplementary material for "Genetic differentiation of water penny beetles may be associated with the formation process of the ancient Lake Biwa": Table S2

**Table S2 COI sequences of CT region used for phylogenic analyses in this study.**

[*1, *2] The species name registered in the database is *E. ramicornis ramicornis* (AB675773 - 779) and *E. ramicornis brunneicornis* (AB675780 - 784), respectively.

[*3] See Table S3 and S4 for detailed information.

References: Hayashi and Sota 2008 (ref1), Hayashi et al. 2012 (ref2), Jung et al. 2020 (ref3).

| Species | Collection site | Sample /Voucher No. | Accession No. | Reference |
|---|---|---|---|---|
| *Eubrianax ramicornis* [*1] | Lake Biwa, Shiga, Japan | *3 | LC572207 - LC572248 | this study |
| | Denpo R., Choshikei, Shodoshima Is., Japan | DR111 | AB675773 | ref2 |
| | Denpo R., Choshikei, Shodoshima Is., Japan | DR113 | AB675774 | ref2 |
| | Taniyama R., Izushi, Hyogo, Japan | DR114 | AB675775 | ref2 |
| | Taniyama R., Izushi, Hyogo, Japan | DR115 | AB675776 | ref2 |
| | Taniyama R., Izushi, Hyogo, Japan | DR116 | AB675777 | ref2 |
| | Taniyama R., Izushi, Hyogo, Japan | DR117 | AB675778 | ref2 |
| | Tsuma, Dogo Is., Oki Is., Japan | DR133 | AB675779 | ref2 |
| | Shakunouchi-koen, Kisuki, Unnan, Shimane, Japan | DR10 | EU287832 | ref1 & 2 |
| | Yoshidomi, Shisuicho, Kikuchi, Kumamoto, Japan | DR83 | EU287833 | ref1 |
| | Imazaike, Okayama, Okayama, Japan | DR20 | EU287834 | ref1 & 2 |
| | Imazaike, Okayama, Okayama | DR41 | EU287835 | ref1 & 2 |
| | R. Sasayamagawa, Murakumo, Sasayama, Hyogo, Japan | DR74 | EU287836 | ref1 |
| | Imazaike, Okayama, Okayama, Japan | DR19 | EU287837 | ref1 & 2 |
| | Shakunouchi-koen, Kisuki, Unnan, Shimane, Japan | DR9 | EU287838 | ref1 & 2 |
| | R. Sasayamagawa, Murakumo, Sasayama, Hyogo, Japan | DR76 | EU287839 | ref1 |
| | Ichinotani, Daisencho, Tottori | DR39 | EU287840 | ref1 & 2 |
| | Ichinotani, Daisencho, Tottori | DR18 | EU287841 | ref1 & 2 |
| | R. Sakaigawa, Rokuonji, Izumo, Shimane, Japan | DR60 | EU287842 | ref1 |
| | Ichinotani, Daisencho, Tottori, Japan | DR17 | EU287843 | ref1 & 2 |
| | R. Hironegawa, Otohara, Sanda, Hyogo, Japan | DR72 | EU287844 | ref1 |
| | Jangan-dong, Seogu, Dejoen, Korea | Eub-001 | KY581625 | ref3 |
| | Junggye-ri, Buan-gun, JB, Korea | Eub-006 | KY581630 | ref3 |
| | Junggye-ri, Buan-gun, JB, Korea | Eub-007 | KY581631 | ref3 |
| | Junggye-ri, Buan-gun, JB, Korea | Eub-008 | KY581632 | ref3 |
| | Gung-ri, Yeoju-gun, GB, Korea | Eub-021 | KY581639 | ref3 |
| | Gung-ri, Yeoju-gun, GB, Korea | Eub-022 | KY581640 | ref3 |
| | Gung-ri, Yeoju-gun, GB, Korea | Eub-023 | KY581641 | ref3 |
| *Eubrianax granicollis* | Lake Biwa, Shiga, Japan | *3 | LC572249 - LC572255 | this study |
| | R. Kandogawa, tachikuekyo, Izumo, Shimane, Japan | DR11 | EU287819 | ref1 |
| | R. Hiikawa, Hinobori, Kisuki, Unnan, Shimane, Japan | DR26 | EU287820 | ref1 & 2 |
| | R. Amidagawa, Daisencho, Tottori, Japan | DR14 | EU287821 | ref1 |
| | Mukorouji, Toyota, Shimonoseki, Yamaguchi, Japan | DR30 | EU287822 | ref1 & 2 |
| | R. Hiikawa, Hinobori, Kisuki, Unnan, Shimane, Japan | DR51 | EU287823 | ref1 |
| | R. Hiikawa, Hinobori, Kisuki, Unnan, Shimane, Japan | DR8 | EU287824 | ref1 & 2 |
| | Nakahara, Okayama, Okayama, Japan | DR35 | EU287825 | ref1 & 2 |
| | Yoshidajima, Kaisei, Kanagawa, Japan | DR37 | EU287826 | ref1 & 2 |
| | Tenjinhara, Kawaramachi, Tottori, Tottori, Japan | DR87 | EU287827 | ref1 & 2 |
| | R. Hiikawa, Hinobori, Kisuki, Unnan, Shimane, Japan | DR53 | EU287828 | ref1 |
| | Tenjinhara, Kawaramachi, Tottori, Tottori, Japan | DR88 | EU287829 | ref1 & 2 |
| | R. Amidagawa, Daisencho, Tottori, Japan | DR13 | EU287830 | ref1 |
| | R. Hiikawa Hinobori, Kisuki Unnan, Shimane, Japan | DR7 | EU287831 | ref1 & 2 |
| | Nakatsu R., Morioka, Iwate, Japan | DR137 | AB675751 | ref2 |
| | Nakatsu R., Morioka, Iwate, Japan | DR139 | AB675752 | ref2 |
| | Warashinagawa, Komukai, Hinata, Aoi, Shizuoka, Japan | DR104 | AB675753 | ref2 |
| | Harada, Dogo Is., Oki Is., Japan | DR129 | AB675754 | ref2 |
| *Eubrianax amamiensis amamiensis* | Kinsakubaru, Amami-oshima Is., Japan | DR093 | AB675746 | ref2 |
| | Kinsakubaru, Amami-oshima Is., Japan | DR094 | AB675747 | ref2 |
| *Eubrianax amamiensis kimurai* | Nago, Okinawa, Japan | DR144 | AB675748 | ref2 |
| | Nago, Okinawa, Japan | DR145 | AB675749 | ref2 |
| | Nago, Okinawa, Japan | DR146 | AB675750 | ref2 |
| *Eubrianax brunneicornis* [*2] | Fukui, Maki, Niigata, Japan | DR118 | AB675780 | ref2 |
| | Fukui, Maki, Niigata, Japan | DR120 | AB675781 | ref2 |
| | Okubogawa, Mizusawaku, Oshu, Iwate, Japan | DR166 | AB675782 | ref2 |
| | Okubogawa, Mizusawaku, Oshu, Iwate, Japan | DR167 | AB675783 | ref2 |
| | Okubogawa, Mizusawaku, Oshu, Iwate, Japan | DR169 | AB675784 | ref2 |
| *Eubrianax ihai* | Hoshino, Ishigaki, Is. , Japan | DR100 | AB675755 | ref2 |
| | Nakamagawa-rindo, Iriomote Is., Japan | DR097 | AB675756 | ref2 |
| | Nakamagawa-rindo, Iriomote Is., Japan | DR099 | AB675757 | ref2 |
| *Eubrianax insularis* | Saruwataridani, Yakushima, Is., Japan | DR023 | AB675758 | ref2 |
| | Saruwataridani, Yakushima, Is., Japan | DR024 | AB675759 | ref2 |
| *Eubrianax lochooensis* | Gima, Kumejima Is., Japan | DR152 | AB675760 | ref2 |
| | Gima, Kumejima Is., Japan | DR153 | AB675761 | ref2 |
| | Gima, Kumejima Is., Japan | DR154 | AB675762 | ref2 |
| | Nago, Okinawa Is., Japan | DR141 | AB675763 | ref2 |
| | Nago, Okinawa Is., Japan | DR142 | AB675764 | ref2 |
| | Nago, Okinawa Is., Japan | DR148 | AB675765 | ref2 |
| | Nago, Is., Japan | DR149 | AB675766 | ref2 |
| *Eubrianax manakikikuse* | Sakita, Taketomi, Iriomote Is., Japan | DR164 | AB675767 | ref2 |
| | Sakita, Taketomi, Iriomote Is., Japan | DR165 | AB675768 | ref2 |
| | Wulai, Taipei, Taiwan | DR003 | AB675769 | ref2 |
| | Wulai, Taipei, Taiwan | DR004 | AB675770 | ref2 |
| *Eubrianax nobuoi* | Niimura, Sumiyo, Amami-oshima Is., Japan | DR201 | AB675771 | ref2 |
| | Niimura, Sumiyo, Amami-oshima Is., Japan | DR202 | AB675772 | ref2 |
| *Eubrianax pellucidus* | Kurokawa R., Kamikurokawa, Kotoura, Tottori, Japan | DR155 | AB675742 | ref2 |
| | Nishiure-toge, Kiyomi-cho, Kamitakara-shi, Gifu | DR161 | AB675743 | ref2 |
| | Takatani, Tsuruoka, Yamagata, Japan | DR105 | AB675744 | ref2 |
| | Kurosonkeikoku, Shimanto, Kochi, Japan | DR107 | AB675745 | ref2 |
| *Eubrianax pellucidus* | Moushi-ohike, Sanda, Hyogo, Japan | DR5 | EU287817 | ref1 & 2 |
| | Moushi-ohike, Sanda, Hyogo, Japan | DR6 | EU287818 | ref1 & 2 |
| *Eubrianax tarokoensis* | Wulai, Taipei, Taiwan | DR001 | AB675785 | ref2 |
| | Wulai, Taipei, Taiwan | DR170 | AB675786 | ref2 |
| *Metaeopsephus japonicus* | Lake Biwa, Shiga, Japan | Bw301 | LC572257 | ref4 |
| *Malacopsephenoides japonicus* | Lake Biwa, Shiga, Japan | Bw311 | LC669411 | ref4 |
