## Supplementary material for "Genetic differentiation of water penny beetles may be associated with the formation process of the ancient Lake Biwa": Table S3

**Psephenidae species collected in this study and their haplotypes or accession numbers for two regions of COI.** Haplotypes are shown for *E. granicollis* and *E. ramicornis* , while accession numbers are shown for other species and Bw199. No haplotype name was assigned for LH of Bw199 and CT of BW197, because both sequences were shorter than other sequences. For accession number for each haplotype, see Table S4. nd: not detected.

| Species name and collection site | Year | Sample ID | Haplotype/ Acc. No. COI (LH) | COI (CT) |
|---|---|---|---|---|
| *Eubrianax granicollis* | | | | |
| Arakawa-ohashi Bridge, Ado River | 2017 | Bw101 | Eg_LH1 | Eg_CT1 |
| | | Bw102 | Eg_LH2 | Eg_CT2 |
| | | Bw103 | Eg_LH3 | Eg_CT3 |
| | | Bw251 | Eg_LH4 | Eg_CT4 |
| Momjibashi Bridge, Echi River | 2017 | Bw197 | Eg_LH1 | LC669412 |
| | | Bw198 | Eg_LH1 | Eg_CT5 |
| | | Bw199 | LC669410 | Eg_CT6 |
| | | Bw254 | Eg_LH1 | Eg_CT7 |
| *Eubrianax ramicornis* | | | | |
| Kaizu | 2017 | Bw026 | Er_LH1 | Er_CT1 |
| | | Bw027 | nd | Er_CT1a |
| | | Bw028 | Er_LH2 | nd |
| | | Bw236 | nd | Er_CT2 |
| | | Bw237 | Er_LH3 | Er_CT1 |
| | | Bw238 | Er_LH2 | Er_CT3 |
| | | Bw239 | Er_LH2 | Er_CT1 |
| | | Bw240 | Er_LH4 | Er_CT2 |
| | | Bw241 | Er_LH2 | Er_CT1 |
| | | Bw242 | nd | Er_CT1 |
| | | Bw243 | Er_LH2 | Er_CT1 |
| | | Bw260 | nd | Er_CT4 |
| | | Bw261 | Er_LH17 | Er_CT5 |
| | | Bw263 | Er_LH18 | Er_CT6 |
| | | Bw264 | Er_LH19 | Er_CT7 |
| | | Bw265 | Er_LH2 | Er_CT8 |
| | | Bw266 | Er_LH2 | Er_CT1 |
| | 2018 | Bw306 | Er_LH16 | nd |
| | | Bw315 | Er_LH2 | Er_CT27 |
| | | Bw316 | Er_LH2 | Er_CT28 |
| | | Bw317 | Er_LH2 | Er_CT1 |
| | | Bw318 | nd | Er_CT5 |

| | | | | |
|---|---|---|---|---|
| | | Bw308 | Er_LH2 | nd |
| | | Bw319 | Er_LH2 | Er_CT3 |
| | | Bw320 | Er_LH1 | Er_CT1 |
| | | Bw321 | Er_LH2 | Er_CT3 |
| | | Bw313 | Er_LH2 | Er_CT1 |
| | | Bw350 | Er_LH2 | Er_CT28 |
| | | Bw351 | Er_LH10 | Er_CT32 |
| | | Bw352 | Er_LH2 | Er_CT1 |
| | | Bw353 | Er_LH2 | Er_CT3 |
| | | Bw354 | Er_LH29 | Er_CT5 |
| Sugaura | 2017 | Bw054 | Er_LH5 | nd |
| | | Bw055 | nd | Er_CT9 |
| | | Bw056 | Er_LH6 | Er_CT9 |
| | | Bw244 | Er_LH7 | Er_CT10 |
| | | Bw245 | Er_LH8 | Er_CT11 |
| | | Bw246 | Er_LH4 | Er_CT12 |
| | | Bw247 | Er_LH9 | Er_CT13 |
| | | Bw248 | nd | Er_CT14 |
| | | Bw249 | nd | Er_CT13 |
| | | Bw250 | Er_LH4 | Er_CT15 |
| | 2018 | Bw307 | Er_LH2 | nd |
| | | Bw322 | Er_LH2 | Er_CT3 |
| | | Bw323 | Er_LH20 | Er_CT9 |
| | | Bw324 | nd | Er_CT29 |
| | | Bw325 | Er_LH21 | Er_CT30 |
| | | Bw326 | Er_LH22 | Er_CT31 |
| | | Bw327 | Er_LH10 | Er_CT32 |
| | | Bw328 | Er_LH23 | Er_CT33 |
| | | Bw329 | Er_LH10 | Er_CT18 |
| | | Bw330 | Er_LH2 | Er_CT1 |
| | | Bw331 | Er_LH24 | Er_CT34 |
| | | Bw332 | Er_LH2 | Er_CT35 |
| | | Bw314 | Er_LH2 | nd |
| | | Bw333 | Er_LH10 | Er_CT41 |
| | | Bw334 | Er_LH30 | Er_CT22 |
| Ukawa | 2017 | Bw149 | Er_LH2 | nd |
| | | Bw150 | Er_LH2 | nd |
| | | Bw151 | Er_LH2 | Er_CT16 |
| Miyagahama | 2017 | Bw161 | Er_LH10 | Er_CT17 |
| | | Bw162 | nd | Er_CT18 |

| | | | | |
|---|---|---|---|---|
| | | Bw163 | Er_LH2 | nd |
| | | Bw252 | Er_LH11 | Er_CT19 |
| | | Bw253 | Er_LH10 | Er_CT17 |
| | 2018 | Bw305 | Er_LH15 | nd |
| | | Bw312 | nd | Er_CT39 |
| | | Bw501 | Er_LH10 | Er_CT17 |
| | | Bw502 | Er_LH10 | Er_CT18 |
| | | Bw503 | Er_LH28 | Er_CT40 |
| | | Bw504 | Er_LH10 | Er_CT18 |
| | | Bw505 | Er_LH10 | Er_CT17 |
| Tsukide | 2018 | Bw302 | Er_LH10 | nd |
| | | Bw335 | Er_LH12 | Er_CT20 |
| | | Bw336 | Er_LH2 | Er_CT21 |
| | | Bw337 | Er_LH10 | Er_CT22 |
| | | Bw338 | Er_LH2 | Er_CT1 |
| | | Bw309 | Er_LH2 | nd |
| Katayama | 2018 | Bw303 | Er_LH2 | nd |
| | | Bw304 | Er_LH13 | nd |
| | | Bw339 | Er_LH2 | Er_CT21 |
| | | Bw340 | Er_LH14 | Er_CT23 |
| | | Bw341 | Er_LH2 | Er_CT21 |
| | | Bw342 | Er_LH13 | Er_CT24 |
| | | Bw343 | Er_LH13 | Er_CT25 |
| | | Bw344 | Er_LH13 | Er_CT26 |
| | | Bw310 | Er_LH25 | nd |
| | | Bw345 | Er_LH2 | Er_CT21 |
| | | Bw346 | Er_LH10 | Er_CT36 |
| | | Bw347 | Er_LH25 | Er_CT37 |
| | | Bw348 | Er_LH26 | nd |
| | | Bw349 | Er_LH27 | Er_CT38 |
| *Mataeopsephus japonicus* | | | | |
| Iso | 2017 | Bw301 | nd | LC572257 |
| *Malacopsephenoides japonicus* | | | | |
| Ado River, estuary | 2017 | Bw078 | LC572256 | nd |
| Katayama | 2018 | Bw311 | LC669409 | LC669411 |
