## Supplementary material for "Genetic differentiation of water penny beetles may be associated with the formation process of the ancient Lake Biwa": Table S4

Table S4 Haplotypes and accession numbers of the two regions of COI.

(a) LH region of *E. ramicornis*

| haplotype | Accession No. |
|---|---|
| Er_LH1 | LC572173 |
| Er_LH2 | LC572174 |
| Er_LH3 | LC572175 |
| Er_LH4 | LC572176 |
| Er_LH5 | LC572177 |
| Er_LH6 | LC572178 |
| Er_LH7 | LC572179 |
| Er_LH8 | LC572180 |
| Er_LH9 | LC572181 |
| Er_LH10 | LC572182 |
| Er_LH11 | LC572183 |
| Er_LH12 | LC572184 |
| Er_LH13 | LC572185 |
| Er_LH14 | LC572186 |
| Er_LH15 | LC572187 |
| Er_LH16 | LC572188 |
| Er_LH17 | LC572189 |
| Er_LH18 | LC572190 |
| Er_LH19 | LC572191 |
| Er_LH20 | LC572192 |
| Er_LH21 | LC572193 |
| Er_LH22 | LC572194 |
| Er_LH23 | LC572193 |
| Er_LH24 | LC572196 |
| Er_LH25 | LC572197 |
| Er_LH26 | LC572198 |
| Er_LH27 | LC572199 |
| Er_LH28 | LC572200 |
| Er_LH29 | LC572201 |
| Er_LH30 | LC572202 |

(b) CT region of *E. ramicornis*

| haplotype | Accession No. |
|---|---|
| Er_CT1 | LC572207 |
| Er_CT1a | LC572208 |
| Er_CT2 | LC572209 |
| Er_CT3 | LC572210 |
| Er_CT4 | LC572211 |
| Er_CT5 | LC572212 |
| Er_CT6 | LC572213 |
| Er_CT7 | LC572214 |
| Er_CT8 | LC572215 |
| Er_CT9 | LC572216 |
| Er_CT10 | LC572217 |
| Er_CT11 | LC572218 |
| Er_CT12 | LC572219 |
| Er_CT13 | LC572220 |
| Er_CT14 | LC572221 |
| Er_CT15 | LC572222 |
| Er_CT16 | LC572223 |
| Er_CT17 | LC572224 |
| Er_CT18 | LC572225 |
| Er_CT19 | LC572226 |
| Er_CT20 | LC572227 |
| Er_CT21 | LC572228 |
| Er_CT22 | LC572229 |
| Er_CT23 | LC572230 |
| Er_CT24 | LC572231 |
| Er_CT25 | LC572232 |
| Er_CT26 | LC572233 |
| Er_CT27 | LC572234 |
| Er_CT28 | LC572235 |
| Er_CT29 | LC572236 |
| Er_CT30 | LC572237 |
| Er_CT31 | LC572238 |
| Er_CT32 | LC572239 |
| Er_CT33 | LC572240 |
| Er_CT34 | LC572241 |
| Er_CT35 | LC572242 |
| Er_CT36 | LC572243 |
| Er_CT37 | LC572244 |
| Er_CT38 | LC572245 |
| Er_CT39 | LC572246 |
| Er_CT40 | LC572247 |
| Er_CT41 | LC572248 |

(c) LH region of *E. granicollis*

| haplotype | Accession No. |
|---|---|
| Eg_LH1 | LC572203 |
| Eg_LH2 | LC572204 |
| Eg_LH3 | LC572205 |
| Eg_LH4 | LC572206 |

(d) CT region of *E. granicollis*

| haplotype | Accession No. |
|---|---|
| Eg_CT1 | LC572249 |
| Eg_CT2 | LC572250 |
| Eg_CT3 | LC572251 |
| Eg_CT4 | LC572252 |
| Eg_CT5 | LC572253 |
| Eg_CT6 | LC572254 |
| Eg_CT7 | LC572255 |
