## Supplementary material for "Genetic differentiation of water penny beetles may be associated with the formation process of the ancient Lake Biwa": Table S1

**Table S1 Anneeling temperature and number of cycles for PCR primers used in this study.**

| target region | Primers | | | Annealing temperature (°C) | no of PCR cycles |
|---|---|---|---|---|---|
| | name | sequence (5' – 3') | reference | | |
| LH | LCO1490 | GGTCAACAAATCATAAAGATATTGG | Folmer et al. 1994 | 51 | 35 |
| | HCO2198 | TAAACTTCAGGGTGACCAAAAAATCA | Folmer et al. 1994 | | |
| CT | COS-Psephenid | CAAGAAAGAGGAAAAAAGGAAAC | Hayashi and Sota 2008 | 55 | 30 |
| | COA-Psephenid | GGGGTTTAARTCCATTGCAC | Hayashi and Sota 2008 | | |
| CT | C1-J-2195 | TTGATTTTTTGGTCATCCAGAAGT | Simon et al. 1994 | 51 | 30 |
| | TL2-N-3014 | TCCAATGCACTAATCTGCCATATTA | Simon et al. 1994 | | |
